# Sulcal Widening in Schizophrenia Maps onto Sulcal Hubs and Energy-Synaptic Genes

**DOI:** 10.64898/2026.01.09.698648

**Authors:** Javier González-Peñas, Hugo G. Schnack, Carmen Rueda Hernández, Covadonga M. Díaz-Caneja, Celia de la Fuente Montero, Marta Martín Echave, Alberto Mora, Niels Janssen, Pedro M. Gordaliza, Alberto Fernández-Pena, Daniel Martín de Blas, Susana Carmona, Wiepke Cahn, Neeltje E.M. van Haren, René S. Kahn, Hilleke Hulshoff Pol, Celso Arango, Yasser Alemán-Gómez, Joost Janssen

## Abstract

Schizophrenia (SZ) is increasingly framed as a disorder of large-scale brain networks emerging from atypical early neurodevelopment, yet how network architecture relates to cortical folding abnormalities remains unclear. Here, we introduce a sulcal morphological-centred network framework that integrates normative modelling of sulcal width with diffusion-derived structural connectivity and transcriptomic data in a large multisite cohort (n = 5,392; 377 SZ). Individuals with SZ showed widespread sulcal widening, affecting 30 of 40 sulci and most pronounced in frontal, temporal and occipital regions. Critically, sulci with higher degree centrality, reflecting greater embedding within the structural connectome, exhibited disproportionately greater widening in SZ (*p*_spin_ = 0.02), indicating that network hubs of cortical folding are preferentially affected. Transcriptomic integration using partial least squares regression identified a single component explaining 56.5% of SZ-related sulcal widening variance (*p*_perm_ = 0.041), implicating genes enriched for synaptic signalling and energy metabolism with adult cortical expression bias and genetic enrichment for cross-disorder psychiatric risk. In contrast, oppositely weighted genes showed prenatal expression bias and enrichment for rare disruptive variants in autism spectrum disorder. Together, these findings link aberrant sulcal morphology in SZ to the brain’s network topology and molecular architecture, positioning cortical folding as a network-embedded phenotype in SZ.

## Introduction

Cortical folding is shaped by a complex interplay of genetic, mechanical and connectivity-driven processes, and has long been hypothesized to reflect constraints imposed by axonal wiring (Van Essen, 1997). Consistent with this view, folding patterns align with cytoarchitecture and connectivity gradients, and emerge early during neurodevelopment (Zilles et al., 2013; Goulas et al., 2014; Sun and Hevner, 2014; Ronan and Fletcher, 2015). In parallel, network neuroscience has demonstrated that highly connected hub regions, compared to less highly connected regions, show disproportionate structural and molecular alterations across psychiatric disorders, including schizophrenia (SZ) (van den Heuvel et al., 2013; Crossley et al., 2014; Wannan et al., 2019; Shafiei et al., 2020; Chopra et al., 2023; Georgiadis et al., 2024). However, these two lines of work have remained largely disconnected: network models have focused on cortical regions rather than folding elements, while studies of sulcal morphology have not been examined within a network-theoretic framework. As a result, it remains unknown whether alterations in cortical folding in SZ preferentially affect sulci that occupy central positions within the brain’s structural connectome. This raises the possibility that sulcal morphological abnormalities in SZ are not randomly distributed, but instead reflect the embedding of sulci within the brain’s network architecture.

Sulcal width reflects morphological properties of both cortical gray matter and underlying white matter and may capture early developmental and mechanical influences related to cortical expansion, tension, and connectivity (Alemán-Gómez et al., 2013; Tallinen et al., 2014; Pizzagalli et al., 2020; Garcia et al., 2025). Atypical sulcal morphology in SZ may reflect disruptions in these prenatal and perinatal processes (Cachia et al., 2015; Garrison et al., 2015; Guo et al., 2015; Palaniyappan et al., 2016). From a genetic perspective, previous studies have estimated the SNP-based heritability of sulcal morphology to range between approximately 15% and 45% (Guen et al., 2019), values comparable to those reported for other cortical morphometric phenotypes (Brouwer et al., 2017; Sun et al., 2022). In a recent genome-wide analysis of different sulcal morphological metrics, more than half of the genome-wide significant loci and the largest heritability estimates were identified for sulcal width compared to other cortical morphometric metrics (Sun et al., 2022). Notably, sulcal width exhibited weaker phenotypic and genetic correlations with other sulcal metrics than those metrics did among themselves, underscoring its biological specificity. Furthermore, sulcal width has been shown to have the strongest genetic correlation with SZ among all sulcal features (Sun et al., 2022). Collectively, these findings position sulcal width as a distinct and genetically informative marker of the early neurodevelopmental mechanisms associated with SZ. However, whether SZ-related alterations in sulcal width are systematically related to large-scale network organization, and whether such relationships are underpinned by specific molecular and developmental programs, remains unknown.

Here, we apply normative modeling to sulcal width data of individuals with SZ to quantify individual deviations from expected trajectories, accounting for demographic, site-related, and image quality-related variability (Marquand et al., 2016; Bethlehem et al., 2022; Di Biase et al., 2022). By combining this approach with a sulci-based connectomic analysis of white-matter connectivity derived from diffusion MRI, we introduce a novel framework to examine whether intrinsic structural brain network organization is associated with individualized sulcal morphological abnormalities in SZ. To probe the molecular mechanisms linking sulcal width and network architecture in SZ, we combined spatial patterns of SZ-related sulcal width alterations with sulci-specific transcriptomic data, situating these changes within the framework of structural brain network organization. Together, this framework enables an investigation of how cortical folding abnormalities in SZ relate to connectomic topology and underlying molecular architecture.

## Subjects and Methods

### Sample characteristics

We analyzed neuroimaging data from 5392 healthy individuals and individuals with SZ, collected across 15 scanning sites. The age distributions per scan site are given in Supplementary Figure 1. The scanner details, sample size, demographics, and clinical characteristics of each scan site, after various exclusions based on data quality, are presented in Supplementary Table 1.

### Image processing and quality control

For a flowchart leading to the final sample, see Supplementary Figure 2. Figure 1 provides a visual summary of the methods used in the current study. We used the same quality-controlled images from the BGS, COBRE, and Utrecht clinical samples as in (Chopra et al., 2023; Martín Echave et al., 2025). Visual quality assessment was part of all quality control procedures for all images from all clinical samples.

**Figure 1.**
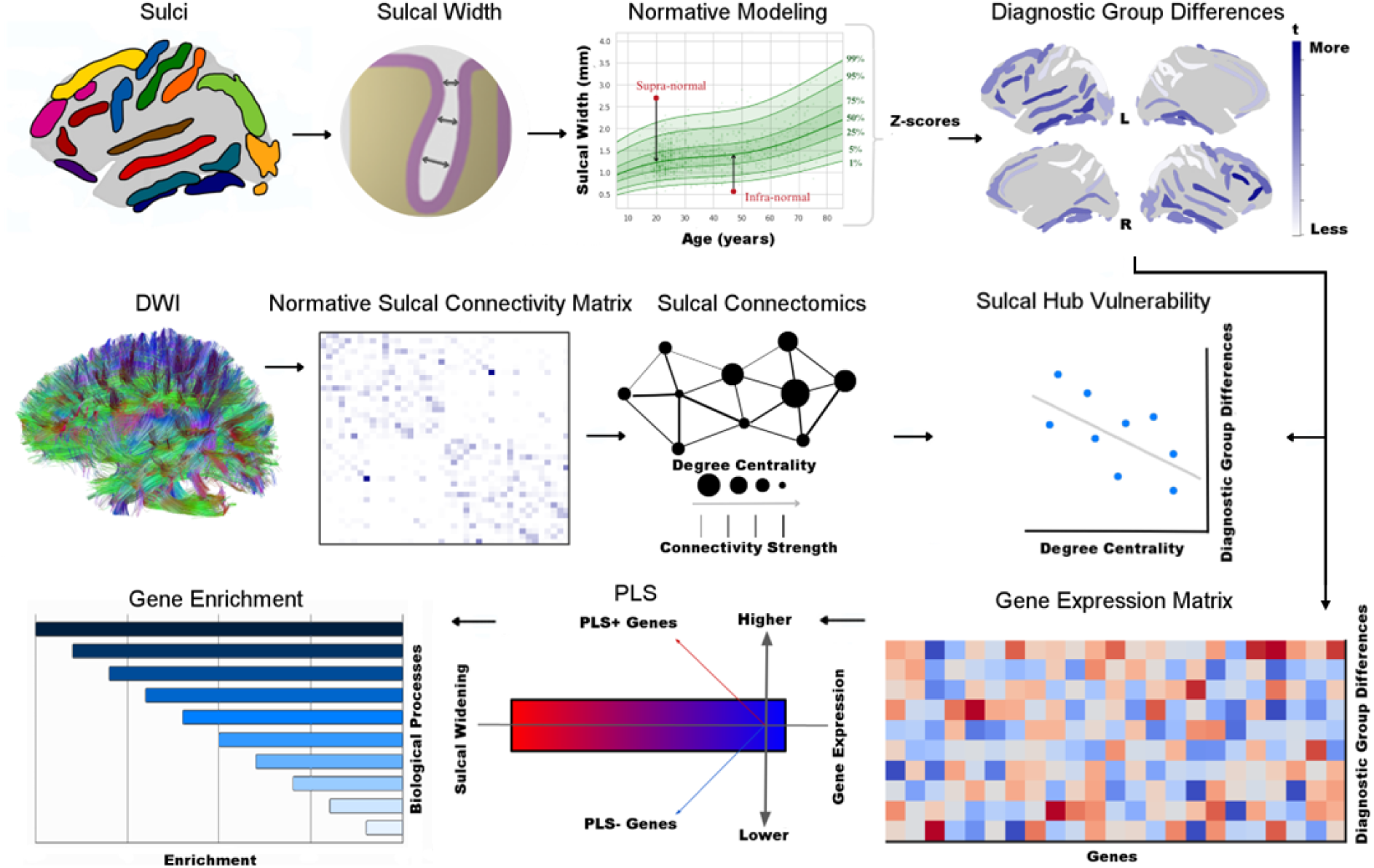
Graphical methods abstract. **Top row left to right:** Sulcal width was measured for 40 bilateral sulci in 5,392 individuals. Normative modeling provided z-scores, representing individual deviations from the normative sulcal width range, which were used in diagnostic group comparisons between individuals with SZ and healthy individuals, producing a vector a t-statistics. **Middle row left to right:** Reference cortico-cortical sulcal connectivity was estimated from diffusion-weighted imaging (DWI) in an independent dataset of 100 healthy individuals. Reference degree centrality was estimated for each sulcus in a first step; in a second step, the reference degree-centrality vector was correlated with the vector of diagnostic group differences (t-statistics). **Bottom row right to left:** Partial least squares regression (PLS) was used to assess the relationship between the vector of t-statistics for the left hemisphere sulci and a sulcal gene expression matrix comprising the expression levels of 15,633 genes for each left hemispheric sulcus. PLS analysis identified genes positively and negatively associated with the t-statistics. Finally, gene enrichment analyses characterized the biological processes and transcriptional underpinnings linked to the diagnostic group differences.

Thereafter, using the included raw T1-weighted images from all datasets, we applied additional image quality control procedures. The Computational Anatomy Toolbox (v12.8.2) was used to generate a weighted overall image quality rating (IQR) for every image (Gaser et al., 2024). IQR combines ratings of basic image properties, including the level of noise and geometric distortions, into a single score that quantifies the overall image quality of a participant’s T1-weighted scan. Lower IQR scores denote higher image quality. As per previous work, we excluded images with an IQR > 2.8 (n=682) (Wolfers et al., 2018). Thereafter, all remaining images were processed centrally using the FreeSurfer analysis suite (v7.1) with default settings (Fischl, 2012). The Freesurfer Euler number was extracted as a proxy for FreeSurfer cortical surface quality (Rosen et al., 2018). Finally, images were removed if the maximum, absolute, within-dataset centered Euler number was larger than 10 (n=5) (Rutherford et al., 2023).

### Sulcal width

For all remaining images, Freesurfer-derived gray and white matter segmentations were processed with BrainVISA (version compiled on August 8th, 2022) using the watershed algorithm for sulcal boundary detection (Mangin et al., 2010). Sulci were extracted using BrainVISA’s Morphologist pipeline, which defines sulci as regions between gyral peaks and skeletonized them to the medial surface between adjacent gyri. We followed the sulcal identification protocol described by Snyder et al. (2024) (Snyder et al., 2024). From 123 potential bilateral sulcal labels, we retained 51. It has been shown that including additional sulci beyond this threshold leads to a significant drop in analyzable data (Snyder et al., 2024). Sulci with inconsistent anatomical relevance (e.g., the anterior lateral fissure, which does not track between gyri) were excluded. Thereafter, using the 51 labeled sulci, adjacent or component sulci were anatomically merged (e.g., combining five pre-central sulcus labels into one; see Supplemental Figure 3 for more details) to address common mislabeling errors by the Morphologist algorithm in spatially proximal regions. This process yielded 20 sulci per hemisphere spanning the lateral, medial, and ventral cortical surfaces (see Supplemental Figure 3), all of which were reliably identified across individuals. Sulcal width was computed for each of the 40 retained sulci using BrainVISA’s Morphologist pipeline. Outlier values—defined as unusually large or small sulcal width values—were identified using the Local Outlier Factor (LOF) method, with 10 neighbours and a contamination parameter of 0.05 (Breunig, Markus M. et al., 2000). We computed a multivariate LOF score for each participant based on all their sulcal width and thickness values. Participants whose LOF score exceeded the 95th percentile were considered outlier participants and excluded from the dataset (n=284), resulting in the final sample size of 5,392 individuals (5,015 healthy individuals, 377 individuals with SZ).

Although sulcal width and cortical thickness are related, they reflect distinct anatomical features (Alemán-Gómez et al., 2013). To quantify the relationship between sulcal width and thickness in our data, we used sulcal cortical thickness values from BrainVISA, defined as the average cortical thickness along the sulcal banks of each sulcus. For each sulcus, we applied linear models to predict sulcal width from sulcal cortical thickness, i.e., the cortical thickness of the sulcal banks, and calculated the proportion of unexplained variance (1 − R^2^) using all individuals.

### Normative modeling

For datasets consisting exclusively of healthy controls, individuals were split into a training set (HC_train_, 90%) and a test set (HC_test_, 10%) using the *createDataPartition* function from the *caret* package in R (Kuhn, 2008). Stratification was applied based on age, sex, and scanner. In clinical datasets, including both healthy controls and individuals with SZ, 80% of healthy controls were assigned to HC_train_, while 20% were allocated to HC_test_. All individuals with SZ were included in the HC_test_ set which thus contained 570 healthy controls and 377 individuals with SZ. Both the training and test sets contained individuals scanned on the same devices, ensuring a “within-site split” (Bayer et al., 2022).

We then applied Bayesian Linear Regression (BLR) (Fraza et al., 2021) to construct normative models of sulcal width for each sulcus. The models were trained on HC_train_ (4,378 healthy controls, 56.7% female; age range: 6–95 years), establishing normative ranges of sulcal width based on age, sex, and Euler number. HC_train_ covered the full age range of the clinical cases (16–67 years). For each sulcus, we quantified individual deviations from the normative prediction using z-scores, computed for each sulcus (*s*) and each individual (*n*) as follows:

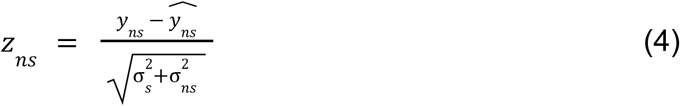

where 𝑦_𝑛𝑠_ is the observed width (computed BrainVisa) and *ŷ*_𝑛𝑠_ is the predicted (normative) width, for sulcus *s* of subject *n*. The difference in these values is normalized to account for two different sources of variation; i) *σ_s_*^2^ which is the aleatoric homoscedastic uncertainty (scalar noise prediction during training) and reflects the variation between individuals across the population, and ii) the epistemic heteroscedastic uncertainty, *σ_ns_*^2^ = *x_ns_^T^*⅀_𝑤_𝑥*_ns_*_’_, where 𝑥*_ns_* is the design vector (i.e., age, sex, Euler number and site) and ⅀_𝑤_ is the posterior covariance of the regression weights (𝑤), which accounts for the variance associated to modeling uncertainty introduced by the model structure or parameter selection (e.g., in age ranges with sparse data).

To validate our model, we applied ten-fold cross-validation within the training cohort. The cohort was divided into ten subsets; in each iteration, BLR models were trained on 90% of the data using age, sex, Euler number, and site as covariates, and evaluated on the remaining 10%. This process was repeated ten times to generate predictions for all individuals in the training set. This approach aligns with standard machine learning practices and provides an unbiased estimate of the model’s generalization performance. Z-scores were calculated and employed to assess the model performance by calculating the mean standardized log-loss, explained variance, standardized mean squared error, root mean squared error, rho, kurtosis, and skewness (see Supplementary Figure 4 for distributions of the evaluation metrics and see Supplementary Figure 5 for the centile plots derived from normative modeling of all 40 sulci, highlighting pronounced nonlinear, monotonically age-related dependency).

To evaluate potential remaining site-related bias in the data, we used linear support vector classifiers (SVCs). Specifically, we trained a series of one-vs-all linear SVCs (with default cost parameter C = 1) to classify scan sites based on normative modeling-based z-scores from healthy controls in both the training and test subsets. For each site, a two-fold cross-validated SVC was used to compute the mean balanced accuracy. Balanced accuracies close to chance level (50%) indicated minimal residual site effects, which was confirmed (see Supplementary Table 2).

### Sulcal structural connectivity matrix generation

One hundred unrelated subjects from the Human Connectome Project (HCP) 1200 Subjects Data Release—referred to as the U100 dataset—were selected to compute the reference sulcal structural connectivity matrix (see the Supplementary Text for a detailed description). Briefly, images were processed using the HCP’s minimal diffusion preprocessing pipeline (Glasser et al., 2013; Sotiropoulos et al., 2013). Multi-shell, multi-tissue constrained spherical deconvolution was applied, yielding white matter fiber orientation distributions. Response functions were estimated using the Dhollander algorithm (Dhollander, T et al., 2016). Deterministic streamline tractography was performed with MRtrix3 and 10 million streamlines were generated for each individual. To enhance the quantitative accuracy of the tractogram, Spherical-deconvolution Informed Filtering of Tractograms (SIFT2) was applied, resulting in weighted streamlines that better reflect the underlying white matter architecture (Smith et al., 2015). These weighted streamlines were then mapped onto the sulcal parcellation (see Supplemental Figure 6 for the sulcal parcellation) to compute the reference sulcus-to-sulcus structural connectivity matrix. From the tractography results, we computed for each subject the number of streamlines connecting each pair of sulci. To reduce the impact of spurious long-distance connections and enhance anatomical plausibility, a connection length-based filtering approach was applied (Betzel et al., 2019). Specifically, a binary mask was derived based on streamline length thresholds to retain only anatomically reliable connections. This mask was then applied uniformly to the reference sulcal structural connectivity matrix. A consistency matrix was also computed where a value of 1 indicates that a particular connection is present in all individual matrices, suggesting a highly reliable and reproducible white matter pathway, while a value of 0 indicates that the connection is never detected across any subject. The sulcal reference weighted degree centrality map was derived from the unthresholded reference sulcus-to-sulcus structural connectivity matrix using the *igraph* R package (Csárdi, G and Nepusz, T, 2005). Reference weighted degree centrality, defined as the sum of all weighted sulcus-to-sulcus connections for each sulcus, was used to identify the degree of ‘hubness’ of a sulcus.

### Sulcal gene expression matrix generation

The abagen toolbox was used to generate a sulcal gene expression matrix from the Allen Human Brain Atlas using the left hemispheric sulcal parcellation in ‘fsaverage’ space (Markello et al., 2021) (see Supplemental Figure 6 for the left hemispheric sulcal parcellation). Processing was conducted using default settings, which include: intensity-based filtering of microarray probes (retaining probes above background intensity in ≥50% of samples), selection of representative probes based on maximum correlation to other probes targeting the same gene, sample-to-region matching using MNI coordinates with a 2 mm tolerance, normalization of expression values within each donor across samples and genes using a scaled robust sigmoid function, and aggregation of expression values within each sulcal basin by averaging across assigned tissue samples. This procedure produced a gene expression matrix where rows correspond to the distinct left hemispheric sulci basins and columns correspond to genes (n = 15,633).

### Statistical analyses

#### Diagnostic group effect on individual deviance per sulcus

For each sulcus, individual z-scores were compared for HC_test_ and SZ groups using Welch t-tests with FDR (*q* < 0.05) as a correction for multiple comparisons and Cohen’s d as a measure of effect size. We conducted a Leave-One-Site-Out (LOSO) analysis to assess the robustness of diagnostic group differences. Normative modeling was repeated for each iteration, excluding one site at a time.

#### Group-level hub vulnerability modeling

To test the sulcal hub vulnerability hypothesis —namely, that sulci with higher reference degree centrality are more susceptible to SZ-related morphometric alterations — we computed the spatial Spearman correlation between the SZ-related morphometric map, i.e., the vector of t-values from the diagnostic group comparisons, and the sulcal reference weighted degree centrality map. To correct for spatial proximity, we used a spatial spin test (Alexander-Bloch et al., 2018). This method involves randomly rotating the sulcal width difference map on the cortical surface, thereby preserving low-order spatial properties—such as spatial autocorrelation—while disrupting the specific spatial associations under investigation. To generate a null distribution, we performed 10,000 surface-based rotations. Statistical significance was determined using a two-tailed *p* threshold of < 0.05. To assess the robustness of the association between the SZ-related morphometric map and sulcal reference degree centrality, we utilized the consistency matrix. Specifically, we applied thresholds to the normative sulcal structural connectivity matrix by retaining only those connections observed in more than 20%, 40%, 60%, or 80% of individuals. For each threshold, we recomputed the spatial correlation between the SZ-related morphometric map and the thresholded reference degree centrality map with significance corrected for by spatial spin tests.

#### Individual-level hub vulnerability modeling

We next examined whether the sulcal hub vulnerability model could be translated to the individual level in SZ. For each individual, we calculated the spatial correlation between their individual deviance profile, i.e. the vector of z-scores from each sulcus, and the sulcal reference degree centrality map. We generated a null distribution of *p*-values using spatial spin tests, in which approximately 2.5% of positive spatial correlations would be expected to reach statistical significance by chance. We then used a *χ²* test to compare the observed proportion of significant positive correlations in the HC_test_ and SZ groups against this expected rate.

#### Transcriptomics of SZ-related sulcal width abnormalities

To characterize the transcriptomic, functional, and genetic architecture underlying SZ-related sulcal width abnormalities, we combined PLS of sulcal gene expression with comprehensive enrichment analyses, transcriptional profiling, and genetic association studies. PLS followed procedures similar to Morgan et al. (2019) and González-Peñas et al. (2024) (Morgan et al., 2019; González-Peñas et al., 2024). The significance of PLS components was evaluated using permutation testing to determine whether the variance explained exceeded chance levels. Gene-wise weights from the PLS model were z-transformed using standard errors estimated via bootstrapping (10,000 resamples) (see the Supplementary Text for a detailed description). *P*-values were computed and corrected for multiple comparisons using FDR. Genes with *p_FDR_* < 0.05 were designated as significantly associated with SZ-related sulcal width abnormalities (hereafter named PLS genes), and classified as PLS⁺ or PLS⁻ based on their positive or negative associations with SZ-related sulcal width abnormalities, respectively (i.e. PLS⁺ and PLS⁻ genes refer to those over- and underexpressed genes in regions with increased widening, and vice versa for regions with decreased widening).

#### Biological signatures of genes associated with SZ-related sulcal width abnormalities

Hierarchical functional clustering and gene-disease associations from DisGeNET database (Piñero et al., 2017) for PLS⁻ and PLS⁺ gene sets was performed using Metascape (Zhou et al., 2019), with the 15,633 brain-expressed genes as the background. Synaptic enrichments were evaluated using hypergeometric tests with the SynGO database (Koopmans et al., 2019). Transcriptional profiling of PLS genes across brain development was conducted using gene expression data from BrainSpan via FUMA (https://fuma.ctglab.nl/) and prenatal versus postnatal biases were evaluated (Hawrylycz et al., 2012; Watanabe et al., 2017). In addition, we investigated the overrepresentation of PLS genes among those specifically expressed genes in (1) tissues from the Genotype-Tissue Expression (GTEx v8) project (GTEx Consortium, 2013) and (2) 39 differentiated cell types identified in single-cell RNA-sequencing data (Zeisel et al., 2018). Tissue- and cell type–specific genes were defined as those within the top 10% of specificity values, following prior approaches (Bryois et al., 2020). Enrichment significance was determined through resampling, comparing observed overlaps to distributions generated from 10,000 random gene sets drawn from the 15,633 background genes (see the Supplementary Text for a more detailed description).

#### Transcriptional profiling of genes associated with SZ-related sulcal widening

We examined whether PLS genes were overrepresented among differentially expressed genes (DEGs) in SZ and other psychiatric disorders by employing a resampling approach, in which the observed overlap was compared to that of 10,000 randomly generated gene lists drawn from the background set of brain-expressed genes (see the Supplementary Text for a more detailed description) (Gandal et al., 2018).

#### Enrichment for common and rare predisposing variation to psychiatric disorders and neuroimaging phenotypes

We further assessed whether the PLS gene sets were enriched for common genetic risk variants associated with SZ and other psychiatric disorders, as well as for neuroimaging phenotypes, using MAGMA v1.10 (de Leeuw et al., 2015). Additionally, we applied logistic regression to test whether PLS genes were overrepresented among genes significantly enriched for rare disruptive variants in SZ (Singh et al., 2022) and in autism and other neurodevelopmental disorders (Fu et al., 2022). Significant findings were validated through 10,000 random permutations of the PLS gene set (details on the GWAS and whole-exome sequencing (WES) datasets are provided in Supplementary Data 2, and see the Supplementary Text for a more detailed description). *P*-values were computed and corrected for multiple comparisons using FDR.

#### Common and independent biological functionalities of SZ-related sulcal width and cortical thickness abnormalities

Finally, we examined whether the biological functions of genes associated with SZ-related sulcal width abnormalities were unique to sulcal width or overlapped with those linked to cortical thickness alterations observed in SZ, such as baseline reductions or accelerated thinning. Specifically, we compared the PLS-derived gene set related to sulcal width with a gene set associated with cortical thickness measures from a prior study in SZ (González-Peñas et al., 2024). Gene set overlap was evaluated by comparing the observed intersection to 10,000 random permutations drawn from the common pool of brain-expressed genes in both studies (see the Supplementary Text for a more detailed description).

## Results

### Diagnostic group effect on individual deviance per sulcus

For each sulcus, normative modeling–based individual z-scores—adjusted for age, sex, image quality, and site—were averaged separately for the HC_test_ and SZ groups. Individuals with SZ showed diffuse sulcal widening relative to HC, particularly in frontal, temporal, and occipital regions, with low to moderate effect sizes (Cohen’s *d* range: 0.01, 0.69) see Figure 2 (Schober et al. 2018). After correction for multiple comparisons, 30 of the 40 sulci exhibited sulcal widening in the SZ group, with no sulci showing decreased width in SZ (see Supplementary Table 3 and Supplementary Figure 7 for group means, individual z-scores, Cohen’s *d* effect size, and significance for each sulcus). Results from the LOSO analysis replicated the main results indicating that the observed diagnostic group differences were not driven by any single site (see Supplementary Figure 8). The supra-normal sulcal map, representing a higher proportion of individuals with supra-normal z-scores (positive z-scores exceeding 1.96, i.e. individuals with significantly greater than normative sulcal width) in the SZ group compared to the HC_test_ group, closely matched the sulcal map described in Figure 1 (see Supplementary Table 5, and Supplementary Figure 9-C, 9-D and 9-E). There were no diagnostic group differences for infra-normal z-scores (negative z-scores below -1.96, i.e. individuals with significantly smaller than normative sulcal width) (see Supplemental Table 4 and Supplementary Figure 9-A and 9-B).

**Figure 2.**
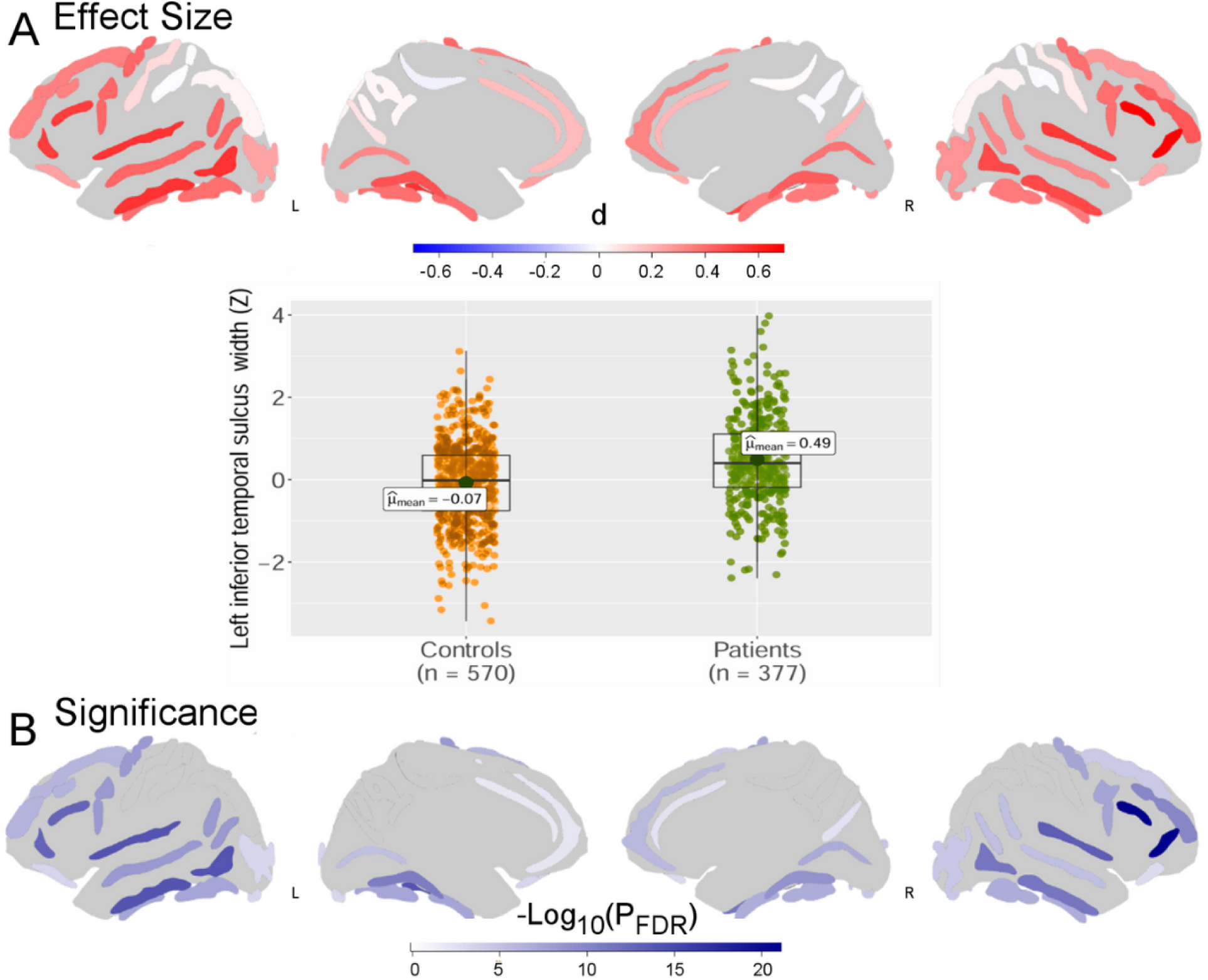
Effect sizes and significance of the diagnostic group differences between HC_test_ and SZ in normative modeling-based z-scores of sulcal width. (**A**) Cohen’s *d* effect sizes for the diagnostic group differences between HC_test_ and SZ in z-scores of sulcal width with a warmer color representing sulcal widening in SZ. Middle panel: Example of individual normative modeling-based z-scores and group means for the left inferior temporal sulcus. (**B**) Log-transformed FDR-corrected *p*-values showing the level of significance for each sulcus (see Supplemental Table 3 and Supplementary Figure 7 for group means, individual z-scores, Cohen’s *d* effect size, and significance for each sulcus).

### Group-level hub vulnerability modeling

The two sulci with the highest reference weighted degree centrality were the left and right paracingulate sulcus (see Supplementary Figure 10 for the complete ranking of sulci by reference weighted degree centrality). To test the hub vulnerability hypothesis in SZ, we compared the spatial distribution of reference weighted degree centrality (see Figure 3-B) with SZ-related increases in sulcal width, i.e. the vector of t-values from the diagnostic group comparisons. There was a positive Spearman correlation between the t-values and degree centrality (⍴ = 0.39, *p*_spin_ = 0.02), indicating that sulcal hubs with higher weighted degree centrality exhibited more pronounced widening in SZ than those with lower weighted degree centrality (see Figure 3-C). This finding replicated at different sparsity-thresholds (see Supplemental Figure 11). This result indicates that sulci with greater sulcus-to-sulcus structural connectivity are more susceptible to SZ-related sulcal widening.

**Figure 3.**
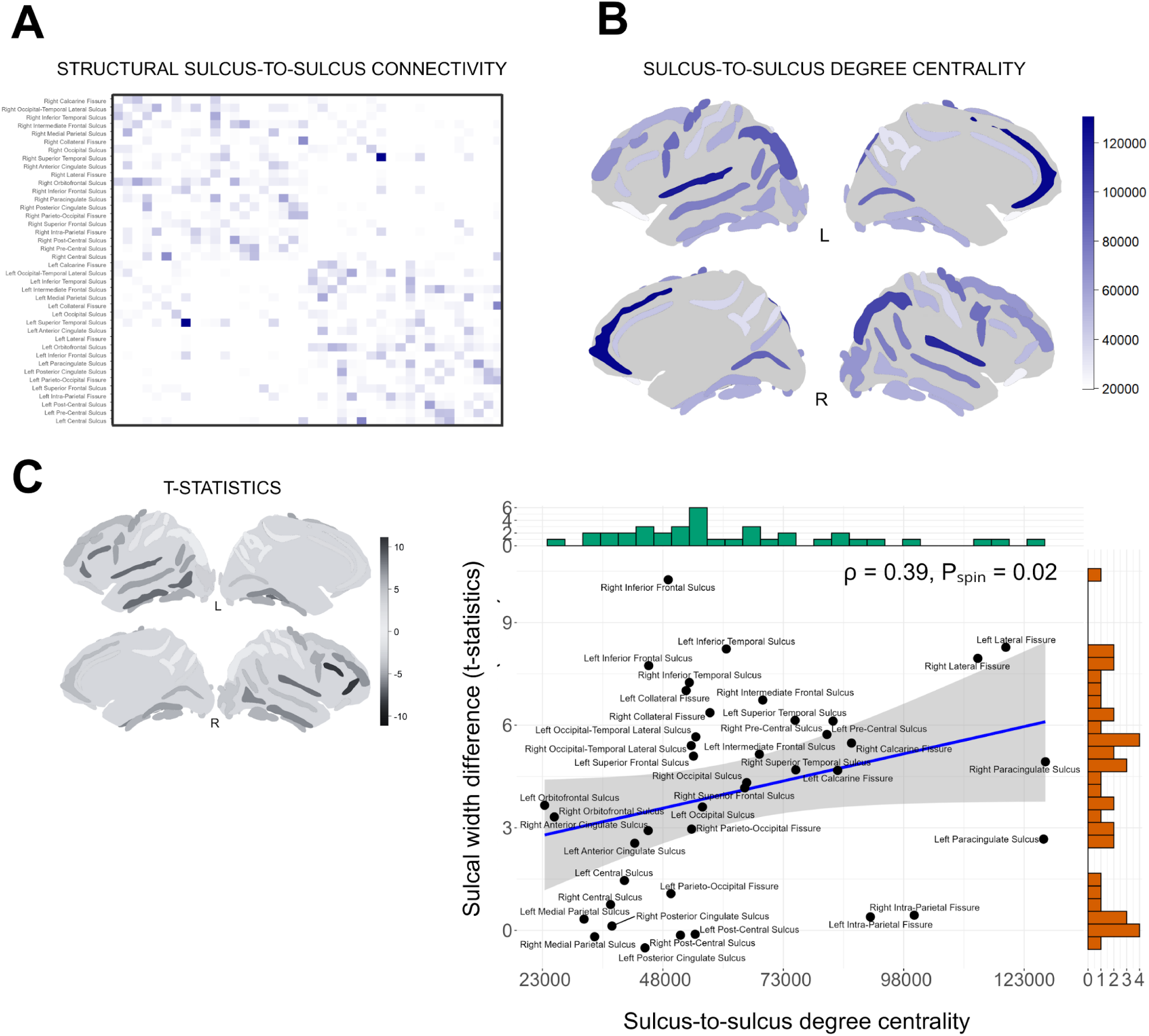
Group-level sulcal hub vulnerability modeling in SZ. (**A**) The reference sulcus-to-sulcus structural connectivity matrix was used to derive reference weighted degree centrality for each sulcus. (**B**) The reference weighted degree centrality map was spatially correlated with the vector of t-statistics from the diagnostic group comparisons (see Supplementary Figure 10 for all sulci ranked by weighted degree centrality). (**C**) There was a positive correlation between the reference weighted degree centrality map and the t-statistics of the diagnostic group differences, indicating that sulci with greater sulcus-to-sulcus structural connectivity tended to show larger SZ-related increases in sulcal width.

### Subject-level hub vulnerability modeling

We tested individual-level hub vulnerability by correlating each subject’s sulcal z-score profile from normative modeling with the reference degree centrality map. Significant positive correlations (*p*_spin_ < 0.05) were found in 6.9% of individuals with SZ, representing a 2.8-fold increase over the 2.5% expected by chance (*χ²* = 28.12, df = 1, *p* < 0.001, see Supplemental Figure 12). In individuals from the HC_test_ group, 3.7% showed a significant positive correlation (*χ²* = 2.81, df = 1, *p* = 0.046) which was significantly lower compared to the individuals with SZ (*χ²* = 4.96, df = 1, *p* = 0.025). This enrichment suggests that individual patterns of sulcal widening are more likely to follow the brain’s sulcal hub architecture than would be expected under the null hypothesis. Individuals with SZ and significant correlations and those without significant correlations did differ from each other in terms of PANSS symptoms or extreme deviations.

### Transcriptomics of SZ-related sulcal widening

We used PLS regression to study the correlation between anatomically patterned brain gene expression from AHBA (15,633 genes) and the SZ-related sulcal widening. A single-component PLS model was retained for further analysis (see Supplemental Text and Supplementary Figures 13 and 14). The first component (PLS1) accounted for 56.51% of the variance in SZ-related sulcal widening, significantly more than expected by chance (*p*_perm_ = 0.041) (see Supplementary Figure 15). After FDR correction, 1,700 genes were significantly associated with SZ-related widening: 1,180 underexpressed (PLS⁻ genes) and 520 overexpressed (PLS⁺ genes) in regions with increased width in SZ (see Supplementary Data 3 for a full list of PLS genes).

### Biological signatures of genes associated with SZ-related sulcal widening

Our functional analysis of the PLS gene sets revealed significant enrichment of the PLS⁻ genes in mitochondrial and energy metabolism pathways (see Figure 4-A), including aerobic respiration (*p*_FDR_ = 8.56 × 10^-9^), ATP synthesis (*p*_FDR_ = 0.02), mitochondrial organization (*p*_FDR_ = 8.42 × 10^-4^), or glycolysis/gluconeogenesis (*p*_FDR_ = 5.22 × 10^-3^), and were also overrepresented in core neuronal system functions (*p*_FDR_ = 0.02; see Supplementary Data 4). In gene-disease association analyses, PLS⁻ genes were linked to metabolic disorders (elevated CSF lactate levels; *p*_FDR_ = 0.002) and neurodegenerative conditions marked by energy dysfunction, such as Leigh disease (*p*_FDR_ = 0.037) and progressive spastic paraplegia (*p*_FDR_ = 0.025; see Supplementary Data 4). Additionally, PLS⁻ genes showed significant enrichment for synaptic processes (see Figure 4-B), including processes at the presynapse (*p*_FDR_ = 8.25 × 10⁻³), synaptic vesicle cycles (*p*_FDR_ = 8.25 × 10⁻³) and transport, presynaptic transmission (*p*_FDR_ = 8.25 × 10⁻³), among others, based on SynGO annotations (see Supplementary Data 5). Together, these findings highlight the central role of mitochondrial dysfunction and synaptic processes in the PLS⁻ gene set. In contrast, no significant functional enrichments were detected for the PLS⁺ genes after FDR correction.

**Figure 4.**
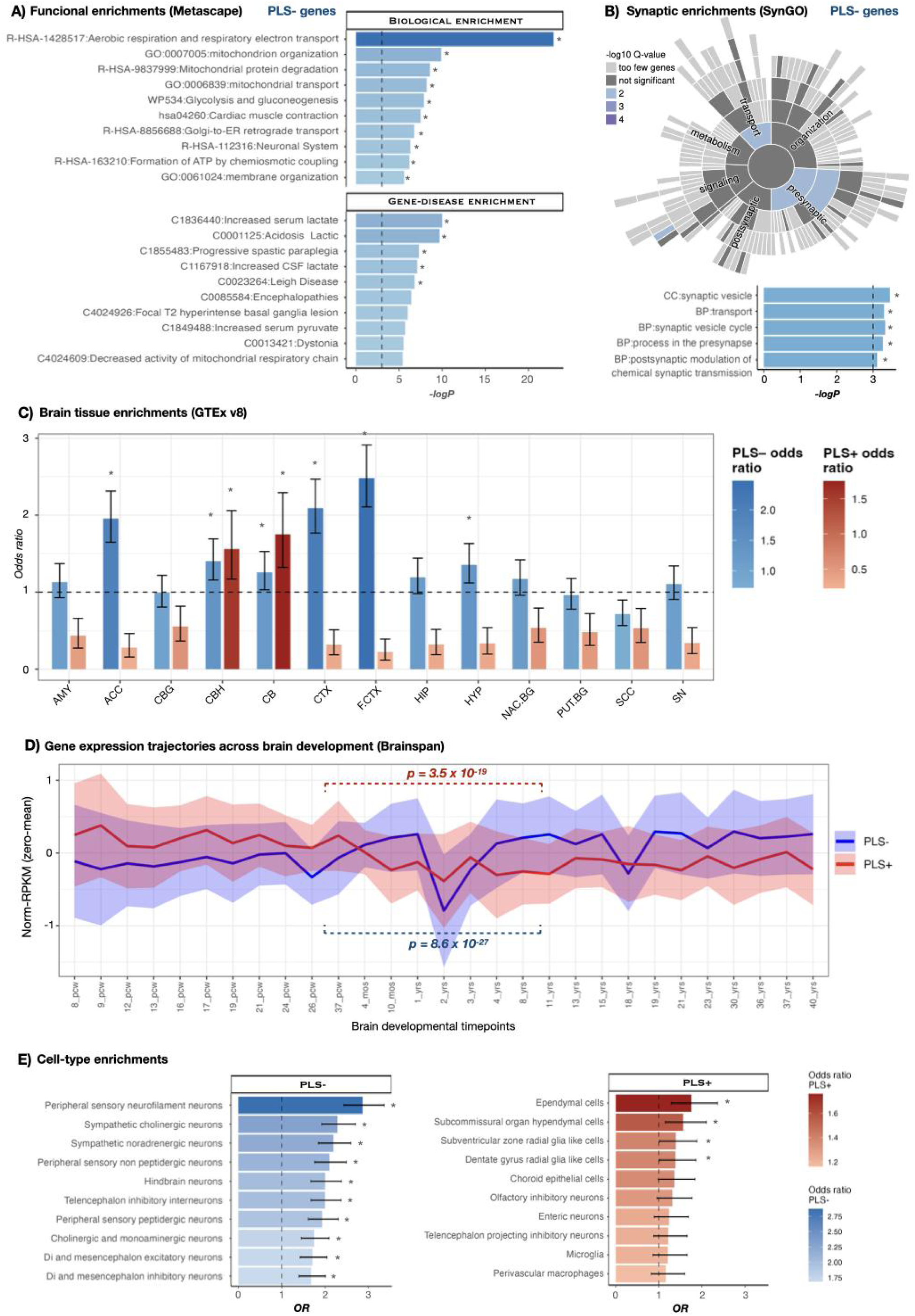
Functional enrichments and transcriptional profiling of genes related with SZ-related sulcal widening. (**A**) Hierarchical functional clustering and gene-disease associations of PLS⁻ genes were performed using custom gene set enrichment analyses in Metascape (Zhou et al., 2019). The 10 most enriched pathways and gene-disease associations are represented. Only PLS⁻ genes had significant enrichments after FDR correction. (**B**) Synaptic enrichment of PLS genes was evaluated using SynGO v1.1 (Koopmans et al., 2019) (https://www.syngoportal.org/) using one-sided Fisher’s exact tests. The five most enriched SynGO terms are represented. Only PLS⁻ genes had significant enrichments after FDR correction. (**C**) Overrepresentation of PLS genes among those specifically expressed across the 13 brain regions from GTEx v8 was assessed by resampling, comparing observed overlaps to 10,000 simulated overlaps. Specifically expressed genes were defined as those in the top 10% of specificity values for each tissue (see Methods). Brain regions with overrepresentation of PLS genes across specifically expressed genes were marked with “*”. Error bars represent OR CI95%. (**D**) Gene expression profiles of PLS⁻ and PLS⁺ genes across human brain development from BrainSpan (http://www.brainspan.org/) (Hawrylycz et al., 2012). Normalized gene expression (RPKM) values per gene were averaged across each PLS gene set at developmental timepoints.The line and shaded contour represent the mean expression and 95%CI for each timepoint. Differences in PLS- and PLS+ gene expression between prenatal and postnatal stages were tested using Wilcoxon rank-sum tests. (**E**) Overrepresentation of PLS genes among those specifically expressed across 39 differentiated cell types from 19 mouse central and peripheral nervous system regions (Zeisel et al., 2018) was assessed by resampling, comparing observed overlaps to 10,000 simulated overlaps. Specific genes were defined as those in the top 10% of specificity values for each cell-type (see Methods). The 10 most enriched cell-types for each PLS gene set are shown. Error bars represent OR CI95%. “*” represents FDR-corrected significant enrichments. AMY = amygdala, ACC = Anterior cingulate cortex, CBG = Caudate basal ganglia, CBH = Cerebellum hemisphere, CB = Cerebellum, CTX = Cortex, F.CTX = Frontal cortex, HIP = hippocampus, HYP = Hypothalamus, NAC.BG = Nucleus accumbens basal ganglia, PUT.BG = Putamen basal ganglia, SCC = Spinal cord cervical c1, SN = Substantia nigra.

### Transcriptional profiling of genes associated with SZ-related sulcal widening

At the adult brain transcriptional level, both PLS⁻ and PLS⁺ gene sets were enriched for genes highly expressed in the cerebellum (PLS⁻: OR = 1.40, 95% CI = 1.16–1.69, *p*_FDR_ = 0.0034; PLS⁺: OR = 1.56, 95% CI = 1.17–2.05, *p*_FDR_ = 0.0055; see Figure 4-C). However, PLS⁻ genes additionally showed strong enrichment in cortical tissue (frontal cortex: OR = 2.47, 95% CI = 2.10–2.91, *p*_FDR_ = 7.9 × 10⁻⁴) and in the hypothalamus (OR = 1.35, 95% CI = 1.12–1.63, *p*_FDR_ = 0.0039). Beyond the brain, PLS⁻ genes were significantly enriched in cardiac and muscle tissues, consistent with their involvement in energy-demanding biological processes (see Supplementary Data 6 and Supplementary Figure 16).

Developmentally, PLS gene sets exhibited distinct expression trajectories: PLS⁻ genes were biased toward postnatal expression (Wilcoxon test, *p* = 8.6 × 10⁻²⁷), whereas PLS⁺ genes were biased toward prenatal expression (Wilcoxon test, *p* = 3.5 × 10⁻¹⁹; see Figure 4-D).

At the cell-type level, PLS⁻ genes were enriched in neuronal populations, including those involved in somatosensory stimulus detection (peripheral sensory neurons: OR = 2.87, 95% CI = 2.44–3.37, *p*_FDR_ = 6.2 × 10⁻⁴; peripheral sensory non-peptidergic neurons: OR = 2.10, 95% CI = 1.76–2.50, *p*_FDR_ = 6.2 × 10⁻⁴), sympathetic nervous system neurons (sympathetic cholinergic: OR = 2.29, 95% CI = 1.93–2.71, *p*_FDR_ = 6.2 × 10⁻⁴; noradrenergic neurons: OR = 2.20, 95% CI = 1.85–2.60, *p*_FDR_ = 6.2 × 10⁻⁴), and hindbrain neurons involved in sensory-motor integration (OR = 2.00, 95% CI = 1.68–2.39, *p*_FDR_ = 6.2 × 10⁻⁴). In contrast, PLS⁺ genes showed modest enrichment in secretory glial and ventricular cell types, including ependymal cells (OR = 2.47, 95% CI = 2.10–2.91, *p*_FDR_ = 7.9 × 10⁻⁴) and subcommissural organ hypendymal cells (OR = 2.47, 95% CI = 2.10–2.91, *p*_FDR_ = 7.9 × 10⁻⁴; see Figure 4-E and Supplementary Data 7).

### Genetic relationship between SZ-related sulcal widening and other psychiatric disorders

We investigated the enrichment of PLS genes among those found to be dysregulated in SZ and other psychiatric disorders based on post-mortem studies, see Figure 5-A. Genes upregulated in SZ showed significant enrichment for PLS⁺ genes (OR = 2.01, 95% CI = 1.67–2.44, *p* = 1 × 10⁻⁵), while genes downregulated in SZ were enriched for PLS⁻ genes (OR = 2.98, 95% CI = 2.62–3.39, *p* = 1 × 10⁻⁵). These findings suggest a shared underlying pathophysiology for sulcal widening and transcriptional dysregulation in SZ. Similar enrichment patterns were seen for genes dysregulated in bipolar disorder (BD) and autism spectrum disorders (ASD), but not for major depression or alcoholism (see Figure 5-A and Supplementary Data 8).

**Figure 5.**
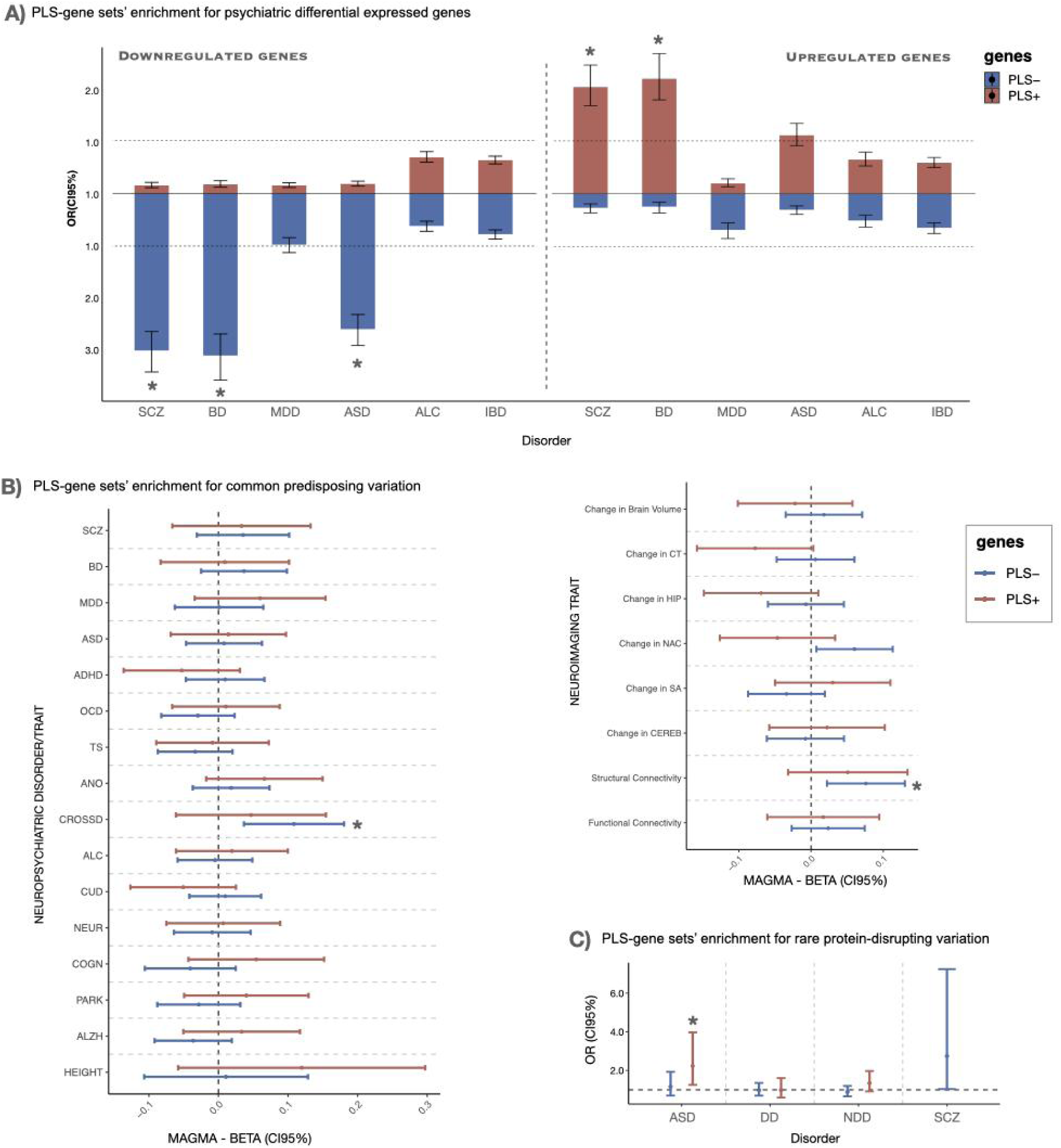
Enrichment of differentially expressed genes (DEG) and predisposing variation to psychiatric disorders across genes associated with SZ-related sulcal widening. (**A**) Overrepresentation of PLS⁺ and PLS⁻ gene sets across genes previously described to be up- and downregulated (p < 0.05) in SZ and other disorders (i.e., bipolar disorder (BD), major depression (MDD), autism spectrum disorders (ASD), alcohol abuse disorder (ALC), and inflammatory bowel disease (IBD) as a non-psychiatric control) (Gandal et al., 2018). “*” represents FDR-corrected significant enrichment after a resampling procedure to test enrichment of PLS gene sets (OR) in DEG genes by comparing against 10,000 randomly selected gene lists selected from background genes (N =15,633). (**B**) Enrichment of PLS⁻ and PLS⁺ genes in predisposing common variation from SZ and other disorders/traits. Enrichment was assessed with MAGMA v1.10 using a one-tailed competitive test, with brain-expressed genes (N=15,633) as background genes. Genetic variation within gene boundaries of 35 kb upstream and 10 kb downstream of the gene bodies was included. Summary data used for analyses is described in Supplementary Data 2. (**C**) Enrichment of PLS⁻ and PLS⁺ genes in rare disruptive variation from SZ and ASD whole exome sequencing (WES) studies. Overrepresentation of PLS genes across 32 SZ-risk genes with *p*_FDR_ < 0.05 (Singh et al., 2022), and 185, 477, and 635 risk genes (TADA-*p_FDR_*< 0.05) for ASD, Developmental Disorder (DD), and Neurodevelopmental Disorder (Fu et al. 2022) (NDD, considering ASD and DD together) was evaluated with logistic regression models using gene length as a covariate. Data used for analyses is described in Supplementary Data 2. Only brain-expressed background genes (N =15,633) were used. “*” represents significant association after permutation procedure. ADHD = Attention deficit and hyperactivity disorder, OCD = obsessive compulsive disorder, TS = Tourette syndrome, ANO = Anorexia nervosa, ALC = Alcohol use disorder, CUD = Cannabis use disorder, NEUR = Neuroticism, COGN = Cognition, PARK = Parkinson disease, ALZH = Alzheimer disease, CROSSD = Psychiatric cross disorder.

Although no significant enrichment of PLS genes was detected for common genetic risk variation associated with SZ or other specific neuropsychiatric disorders, PLS⁻ genes were significantly enriched for the cross-disorder psychiatric genetic factor (BETA = 0.11, 95% CI = 0.04–0.18, *p*_FDR_ = 0.046; see Figure 5-B). In addition, PLS⁻ genes showed significant enrichment for genetic variation associated with structural brain connectivity (BETA = 0.08, 95% CI = 0.02–0.13, *p_FDR_* = 0.046; see Figure 5-B and Supplementary Data 9), consistent with the relationship we report between sulcal widening and sulcal hub vulnerability. Conversely, in relation to rare risk genetic variation, PLS⁺ genes were enriched for genes harboring protein truncating variants for ASD (11 PLS⁺ genes out of 185 ASD genes; OR(CI95%) = 2.23 (1.26;3.97); *p_perm_* = 0.009; see Figure 5-C; Supplementary Data 10).

### Independent biological functions of SZ-related sulcal width compared to those underlying cortical thickness abnormalities

To quantify the relationship between sulcal width and cortical thickness, we fitted linear models predicting sulcal width from sulcal cortical thickness across all individuals and computed the proportion of unexplained variance (1 − R^2^). More than 64% of the variance in sulcal width remained unexplained for all sulci, indicating that variation in sulcal width in our sample is not solely attributable to changes in sulcal cortical thickness (see Supplementary Figure 17).

Finally, we evaluated whether the biological functions associated with the sulcal widening identified in this study are distinct from those linked to cortical thickness alterations in SZ previously reported by our group. Comparing PLS-derived gene sets across studies, we observed a significant overlap (p_perm = 1 × 10⁻⁴; Supplementary Data 11) between PLS⁻ genes associated with sulcal widening and those related to accelerated cortical thinning (ACT-PLS⁻)—that is, genes underexpressed in regions showing both sulcal widening and accelerated cortical thinning in schizophrenia—. No significant overlap with PLS genes related to cortical thickness differences (CCD) was observed. To assess whether the observed PLS⁻ enrichments related to sulcal width abnormalities were driven by this overlapping subset of genes, we repeated all major enrichment analyses on the remaining 934 non-overlapping PLS⁻ genes. Biological and transcriptional profiles were preserved after removing the overlapping genes (Supplementary Text, Supplementary Data 12–16), supporting the independence of the biological underpinnings of sulcal widening reported here.

## Discussion

This study investigates deviations in sulcal width in schizophrenia (SZ) through normative modeling, brain connectivity mapping, and transcriptomic integration. We identified a widespread pattern of sulcal widening across cortical regions, without evidence of sulcal narrowing. The pattern of SZ-associated increased sulcal width followed the centrality principle of brain network organization and correlated with cortical gene expression. The genes driving this association span pre- and postnatal neurodevelopment, are implicated in mitochondrial dysfunction and synaptic pathways, are dysregulated in SZ and related disorders, and are enriched for common risk variation for SZ and rare protein-truncating variants linked to ASD.

Sulcal width indexes the spacing between gyral banks, integrating cortical thinning, subarachnoid expansion, and white-matter changes (Kochunov et al., 2005; Im et al., 2008; Alemán-Gómez et al., 2013; Lin et al., 2025), each of which has been found to be consistently affected in SZ (van Haren et al., 2011; Zhang et al., 2015; Kochunov et al., 2016; Kelly et al., 2018; van Erp et al., 2018; García-San-Martín et al., 2025b). Our findings of widespread increased sulcal width without regional narrowing suggest a one-sided developmental trajectory, reflecting early disruptions in cortical folding dynamics leading to widening (Jin et al., 2018). Mechanically, cortical folding arises from tension between cortical growth and subcortical constraints (Van Essen, 1997; Tallinen et al., 2014), where gyral crowns experience tensile expansion and sulcal fundi undergo compression. Disruption of this equilibrium through altered axonal tension or connectivity could lead to disproportionate sulcal width (Llinares-Benadero and Borrell, 2019). Supporting this interpretation, SZ shows large-scale dysconnectivity and reduced structural coupling between gyral and sulcal regions (Palaniyappan et al., 2011; van den Heuvel et al., 2013; Lin et al., 2025). Sulci with higher network centrality—those acting as structural hubs—displayed greater width, supporting the hub vulnerability hypothesis. Prior studies show that highly connected cortical hubs, defined by their central position in functional and structural networks, exhibit the strongest thickness and deformation abnormalities in schizophrenia (Wannan et al., 2019; Shafiei et al., 2020; Georgiadis et al., 2024). Extending this principle to sulcal morphology, our results indicate that topologically central sulci tend to be more affected, suggesting that cortical folding adheres to the brain’s network and energetic constraints (Llinares-Benadero and Borrell, 2019). Such hub sulci are metabolically costly, relying on dense synaptic and vascular activity, and are therefore particularly vulnerable to oxidative stress, mitochondrial inefficiency, and neuroinflammation—processes repeatedly implicated in SZ (Stephan et al., 2009; Fillman et al., 2013; Heckers and Konradi, 2015; Friston et al., 2016). Using individual-level deviation data from normative modelling, nearly 7% of individuals with SZ showed significantly above-chance correlations between sulcal width deviations from normality and reference network centrality. This modest percentage emphasizes inter-individual heterogeneity in sulcal morphology which is consistent with prior normative modeling work in SZ (Wolfers et al., 2018; Di Biase et al., 2022; Segal et al., 2023; García-San-Martín et al., 2025a). Altogether, these findings indicate that the network organization guiding cortical alterations in SZ extends to sulcal morphology, linking cortical folding abnormalities with intrinsic connectomic and metabolic architecture and emphasizing substantial inter-individual variability in how these network-level vulnerabilities manifest across individuals with SZ.

Our transcriptomic analysis provides molecular validation for the metabolic hub vulnerability framework. The first PLS component (PLS1) explained 56.51% of the variance in the spatial pattern of increased sulcal width in SZ, substantially more than prior PLS-based analyses of cortical thickness phenotypes and thus captures a robust biological component underlying these SZ-related geometric abnormalities (Romero-Garcia et al., 2019; González-Peñas et al., 2024). Negatively weighted genes (PLS-) were markedly enriched for mitochondrial organization, oxidative phosphorylation, and synaptic vesicle cycling. This convergence confirms that the sulci with most width tend to be those with transcriptional downregulation of pathways supporting energy metabolism and synaptic maintenance. Positively weighted genes (PLS+) showed enrichment for glial and developmental programs with prenatal expression peaks, suggesting complementary contributions of early neurodevelopmental factors and later energy-dependent maintenance processes. Together, these transcriptomic findings corroborate the notion that increased sulcal width in SZ arises from the interplay of highly central cortical regions that impose elevated postnatal metabolic demands on neuronal circuits and prenatal cerebellar and glial developmental programs that shape early circuit architecture and vulnerability, jointly increasing susceptibility to later structural degradation.

The PLS- and PLS+ gene sets were respectively downregulated and upregulated in SZ and bipolar disorder, and in autism spectrum disorder (significantly for PLS+), but not in major depressive disorder. Despite the strong transcriptomic overlap between SZ and depression (Gandal et al., 2018), this dissociation indicates molecular specificity linked to increased sulcal width. Although gene sets associated with increased sulcal width and cortical thinning in SZ partially overlapped, the sulcal-related signature remained significantly enriched after removing shared genes, demonstrating distinct underlying mechanisms. Population-genetic analyses further show that sulcal width exhibits phenotypic and genetic correlation structures different from other cortical metrics (Sun et al., 2022), and evidence from neurodegenerative research supports increased sulcal width as a more sensitive indicator of structural disorganization than cortical thickness (Im et al., 2008; Cai et al., 2017; Wang et al., 2021).

At the level of genetic variation, PLS-genes were enriched for common cross-disorder risk but not for SZ-specific variants, indicating that increased sulcal width reflects a shared neurodevelopmental and metabolic vulnerability rather than disorder-specific effects. PLS-genes were also enriched for variation associated with brain structural connectivity, reinforcing the observed correlations between SZ-associated increased sulcal width and network centrality. In contrast, the PLS+ gene set, characterized by a strong prenatal expression bias, was enriched for autism spectrum disorder risk genes carrying rare protein-truncating variants, consistent with mid-fetal cortical developmental programs (Satterstrom et al., 2020; Dear et al., 2024). These rare coding risk genes overlap substantially between ASD and SZ (Fu et al., 2022), aligning with genetic evidence implicating early neurodevelopmental perturbations in cortical architecture and SZ risk.

Together, the imaging and transcriptomic data support a mechanistic model in which increased sulcal width in SZ may arise from failure to maintain the mechanical balance that shapes sulcal morphology. Highly connected hub sulci, which sustain dense synaptic signaling and high metabolic demand, could rely on mitochondrial and synaptic efficiency to preserve axonal tension across the cortical sheet. When energetic or synaptic maintenance processes are compromised, axonal tension may weaken, possibly reducing the compressive forces that stabilize sulcal walls. This mechanical relaxation may produce gradual sulcal expansion and loss of curvature, mirroring the cortical thinning and reduced gyrification consistently observed in SZ (Palaniyappan and Liddle, 2012).

Several limitations should be noted. Our cross-sectional design precludes direct inference about progression, and longitudinal work is needed to determine whether sulcal widening represents a deviation from early neurodevelopmental processes or progressive change. Medication exposure and chronic illness effects, among other epiphenomena, could contribute to variability, though prior studies suggest limited impact on large-scale morphology (van Erp et al., 2018). Transcriptomic mapping from the Allen Human Brain Atlas, while informative, is based on a limited donor sample and may not capture individual variability. Moreover, transcriptomic analyses based on postmortem brain tissue warrant caution (Hernandez et al., 2021). Gene expression varies across the lifespan (Viñuela et al., 2018) and can be influenced by illness course and treatment exposure (Ota et al., 2019). Furthermore, the fewer positive results in the enrichment analyses for PLS+ gene sets, compared with PLS−, may partly reflect reduced statistical power due to the smaller number of PLS+ genes. Finally, while sulcal hubs were defined from group-level networks, individual connectivity patterns may differ (Mueller et al., 2013).

In summary, we show that SZ involves widespread sulcal widening organized along the brain’s network topology and coupled to transcriptional gradients related to mitochondrial and synaptic function. These findings extend the concept of hub vulnerability from cortical thickness and volume to the domain of sulcal width, suggesting that energetic and network constraints may jointly shape the structural phenotype of SZ. Sulcal width thus offers a novel geometric lens on the neurodevelopmental and bioenergetic mechanisms underlying cortical disorganization in SZ.

## Acknowledgements

Supported by the Spanish Ministry of Science and Innovation, Instituto de Salud Carlos III (ISCIII), CIBER-Consorcio Centro de Investigación Biomédica en Red-(CB/07/09/0023), co-financed by the European Union, ERDF Funds from the European Commission, “A way of making Europe”, (PI16/02012, PI17/01249, PI17/00997, PI19/01024, PI20/00721, PI22/01824, PI22/01621, PI23/00625), financed by the European Union - NextGenerationEU (PMP21/00051), Madrid Regional Government (S2022/BMD-7216 AGES 3-CM), European Union Seventh Framework Program, European Union H2020 Program under the Innovative Medicines Initiative 2 Joint Undertaking: Project PRISM-2 (Grant agreement No.101034377), Project COllaborative Network for European Clinical Trials For Children “c4c” (Grant agreement No 777389) Horizon Europe, the National Institute of Mental Health of the National Institutes of Health under Award Number 1U01MH124639-01 (Project ProNET), Award Number 5P50MH115846-03 (Project FEP-CAUSAL) and Award Number 1R01MH128971-01A1 (Project SZ-aging), Fundación Familia Alonso, and Fundación Alicia Koplowitz.

## Ethics approval and consent to participate

All methods were performed in accordance with the relevant guidelines and regulations. All sites obtained local institutional review board approval. Written informed consent was obtained from every participant, or from the participant’s guardian for minors. All studies were conducted in accordance with the Declaration of Helsinki.

## Author contribution

Substantial contributions to the conception or design of the work; or the acquisition, analysis, or interpretation of data for the work: Joost Janssen, Javier González-Peñas, Hugo G. Schnack, Carmen Rueda Hernández and Yasser Alemán-Gómez. Drafting the work or revising it critically for important intellectual content: Javier González-Peñas, Hugo G. Schnack, Carmen Rueda Hernández, Covadonga M. Díaz-Caneja, Celia de la Fuente Montero, Marta Martín Echave, Alberto Mora, Niels Janssen, Pedro M. Gordaliza, Alberto Fernández-Pena, Daniel Martín de Blas, Susana Carmona, Wiepke Cahn, Neeltje E.M. van Haren, René S. Kahn, Hilleke Hulshoff Pol, Celso Arango, Yasser Alemán-Gómez and Joost Janssen. Agreement to be accountable for all aspects of the work in ensuring that questions related to the accuracy or integrity of any part of the work are appropriately investigated and resolved: Javier González-Peñas, Yasser Alemán-Gómez and Joost Janssen.

## Data availability

In addition to non-publicly available datasets, the following publicly available datasets were used: Aomic (id1000, piop1 and piop2) available at https://openneuro.org/datasets/ds003097, https://openneuro.org/datasets/ds002785 and https://openneuro.org/datasets/ds002790; camcan available at https://camcan-archive.mrc-cbu.cam.ac.uk; dlbs available at https://fcon_1000.projects.nitrc.org/indi/retro/dlbs; ixi available at http://brain-development.org/ixidataset; narratives available at https://openneuro.org/datasets/ds002345; oasis3 available at https://www.oasis-brains.org; rockland available at http://fcon_1000.projects.nitrc.org/indi/enhanced; sald available at http://fcon_1000.projects.nitrc.org/indi/retro/sald; BGS available at http://schizconnect.org/; COBRE available at http://schizconnect.org/; LA5C available at https://openneuro.org/datasets/ds000030/versions/00016; SRPBS available at https://bicr-resource.atr.jp/srpbsopen/.

## Code availability

The code will be made available upon acceptance of the manuscript for publication.

