## Supplementary Material for "Sulcal Widening in Schizophrenia Maps onto Sulcal Hubs and Energy-Synaptic Genes"

Running title: Sulcal Widening in Schizophrenia

Javier González-Peñas PhD, Hugo G. Schnack PhD, Carmen Rueda Hernández MS, Covadonga M. Díaz-Caneja MD PhD, Celia de la Fuente Montero MS, Marta Martín Echave MS, Alberto Mora MS, Niels Janssen PhD, Pedro M. Gordaliza PhD, Alberto Fernández-Pena PhD, Daniel Martín de Blas PhD, Susana Carmona PhD, Wiepke Cahn MD PhD, Neeltje E.M. van Haren PhD, René S. Kahn MD PhD, Hilleke Hulshoff Pol PhD, Celso Arango MD PhD, Yasser Alemán-Gómez PhD & Joost Janssen PhD

### Content

#### Supplementary Text: Methods and Results

Normative sulcal structural connectivity matrix generation

Transcriptomics of SZ-related sulcal width abnormalities

Biological signatures of genes associated with SZ-related sulcal width abnormalities

Genetic relationship between SZ-related sulcal width abnormalities and other related disorders

Enrichment for Common predisposing variation to SZ-related disorders and neuroimaging phenotypes

Common and independent biological functionalities of SZ-related sulcal width and cortical thickness abnormalities

#### Supplementary Tables

Supplementary Table 1 is part of the Supplementary Data

Supplementary Table 2

Supplementary Table 3

Supplementary Table 4

Supplementary Table 5

#### Supplementary Figures

Supplementary Figure 1: Age distributions per dataset

Supplementary Figure 2: Flow diagram for inclusion/exclusion

Supplementary Figure 3: Sulcal nomenclature

Supplementary Figure 4: Distributions of evaluation metrics of normative modelling

Supplementary Figure 5: Centile plots of normative modelling

Supplementary Figure 6: Sulcal parcellation in 'fsaverage' space

Supplementary Figure 7: Individual z-scores grouped by diagnosis for each sulcus

Supplementary Figure 8: Leave-One-Site-Out analysis

Supplementary Figure 9: Infra- and supra-normal z-scores comparison

Supplementary Figure 10: Ranking of sulci by weighted degree centrality

Supplementary Figure 11: Structural connectivity matrices at multiple connectivity thresholds

Supplementary Figure 12: Distribution of correlation coefficients

Supplementary Figure 13: RMSEP versus number of PLS components.

Supplementary Figure 14: Variance explained by the first 10 PLS components.

Supplementary Figure 15: Significance of variance explained by a PLS model of one component (PLS1)

Supplementary Figure 16: Overrepresentation of PLS genes among those specifically expressed across the 24 non-brain tissues from GTEx v8

Supplementary Figure 17: Relationship between sulcal width and thickness

References

Normative sulcal structural connectivity matrix generation

*Sample*

One hundred unrelated subjects from the Human Connectome Project (HCP) 1200 Subjects Data Release—referred to as the U100 dataset—were selected to compute the sulcal-based structural connectivity matrix. The U100 subset is widely used in structural and functional connectomics due to its high-quality, multi-shell diffusion imaging, standardized preprocessing pipelines, and demographic diversity while minimizing genetic relatedness (Van Essen et al., 2013). This makes it especially well-suited for population-level analyses of brain connectivity without confounding familial or twin effects. All the details about the acquisition protocols are described in Van Essen et al., 2013. Only the T1-weighted (T1w) and diffusion-weighted (DW) images were downloaded and utilized from the HCP dataset, as these modalities provided the necessary structural and diffusion information for surface reconstruction and connectivity analysis.

*Sulcal parcellation*

We used a cortical parcellation of sulcal banks and fundi in FreeSurfer ‘fsaverage’ space which was consistent with the sulci labelling used in the current study, BrainVISA standard sulcal labeling and endpoints of sulcal regions in the Destrieux atlas (see Supplementary Figure 6 and (Snyder et al., 2024)). The T1w images from the u100 HCP dataset were processed using FreeSurfer v7.4.0 to obtain the cortical surfaces and generate the subject-specific mappings to the ‘fsaverage’ surface space. These surface-based transformations were used to project the sulcal parcellation template onto each individual’s cortical anatomy. In order to get tractography termination masks suitable for the sulcal connectomes' construction, the labeled sulcal regions were dilated towards the subcortical white matter with a radius of 1 mm.

*Sulcal structural connectivity matrix generation*

Alongside the native diffusion-weighted (DW) images, the HCP database also provided preprocessed DW data. These images were processed using the HCP's minimal diffusion preprocessing pipeline, which includes gradient nonlinearity correction, EPI distortion correction, eddy current and motion correction with outlier replacement, and registration to the structural T1-weighted image in AC-PC space (Glasser et al., 2013; Sotiropoulos et al., 2013). For estimating fiber orientation distributions (fODs), multi-shell, multi-tissue constrained spherical deconvolution (MSMT-CSD) was applied using the dwi2fod command, yielding white matter fODs. Response functions were estimated using the Dhollander algorithm (Dhollander, T et al., 2016). Deterministic streamline tractography was performed with MRtrix3's tckgen, employing the SD_STREAM algorithm with anatomically constrained tractography (ACT) priors. Tracking parameters included a step size of 0.5 mm, a minimum streamline length of 5 mm, and a maximum length of 250 mm. For each subject, 10 million streamlines were generated, from which structural connectivity matrices were derived according to the cortical parcellation. To enhance the quantitative accuracy of the tractogram, SIFT2 (Spherical-deconvolution Informed Filtering of Tractograms; (Smith et al., 2015)) was applied, resulting in weighted streamlines that better reflect the underlying white matter architecture. These weighted streamlines were then mapped onto a sulcal parcellation to compute the structural connectivity matrices between the sulci. From the tractography results, we computed for each subject the number of streamlines connecting each pair of sulci. To reduce the impact of spurious long-distance connections and enhance anatomical plausibility, a connection length-based filtering approach was applied following the method proposed by Betzel et al. (2019) (Betzel et al., 2019). Specifically, a binary mask was derived based on streamline length thresholds to retain only anatomically reliable connections. This mask was then applied uniformly to the sulcal connectivity matrix.

Transcriptomics of SZ-related sulcal width abnormalities

The correlation between the brain gene expression matrix (15,633 genes × 20 left hemispheric sulci, derived using the Abagen toolbox, see Markello et al. 2021) and SZ-related morphometric patterns was evaluated using partial least squares regression (PLSR), following approaches similar to those of Morgan et al. (2019) and González-Peñas et al. (2024) (Morgan et al., 2019; González-Peñas et al., 2024). We first determined the number of latent components required to capture the covariance between gene expression and SZ-related sulcal width abnormalities by fitting PLSR models with one to ten components and assessing predictive performance via leave-one-out cross-validation. Based on the root mean squared error of prediction (RMSEP) curve and cumulative variance explained, we found that while two components minimized RMSEP, the first component alone accounted for over 50% of the variance in SZ-related sulcal width abnormalities. Therefore, we retained a single-component model for all downstream analyses.

To assess the statistical robustness of PLS1 and control for false positives, we tested whether the variance explained by PLS1 exceeded chance levels. This was done by generating 10,000 random permutations of the SZ-related sulcal width abnormality values, refitting the one-component PLSR for each permuted dataset, and comparing the observed variance explained to the null distribution. The empirical *p*-value was computed as the proportion of permutations with variance equal to or greater than that observed in the true data.

Next, to characterize individual gene contributions to the correlation between spatial gene expression and SZ-related sulcal width abnormalities, we extracted gene-wise regression coefficients (β). We estimated the standard error (SE) of each β using 10,000 bootstrap resamples of the 20 sulci (sampling with replacement), refitting the PLSR model on each replicate. From these SEs, we calculated two-tailed z-scores and corresponding parametric p-values, which were corrected for multiple testing across 15,633 genes using the Benjamini–Hochberg False Discovery Rate (FDR) procedure (Benjamini and Hochberg, 1995). Genes with $p_{FDR} < 0.05$ were designated as significantly associated with SZ-related sulcal width abnormalities (PLS genes).

Genes were classified as $PLS^+$ or $PLS^-$ based on the sign of their PLS1 weight, indicating positive or negative associations with SZ-related sulcal width abnormalities, respectively. Positive values corresponded to regions showing greater sulcal widening in SZ. Thus, $PLS^+$ genes are those overexpressed and $PLS^-$ genes those underexpressed in regions with increased widening. This analytical pipeline combines cross-validated model selection, permutation testing, and bootstrap inference to identify robust gene–atrophy relationships.

Biological signatures of genes associated with SZ-related sulcal width abnormalities

Hierarchical functional clustering of $PLS^+$ and $PLS^-$ genes was performed using custom gene set enrichment analyses in Metascape (Zhou et al., 2019), drawing on multiple databases, including KEGG Pathway, GO Biological Processes, Reactome Gene Sets, Canonical Pathways, CORUM, and WikiPathways. Enrichment p-values were computed using the cumulative hypergeometric distribution, using FDR to correct for multiple testing ($q < 0.01$). Significant terms were hierarchically clustered into a tree structure based on Kappa-statistical similarities in gene membership (kappa score ≥ 0.3).

We additionally assessed significant gene-disease associations using the DisGeNET database (Piñero et al., 2017). Synaptic enrichment of PLS genes was evaluated using SynGO v1.1 (https://www.syngoportal.org/) (Koopmans et al., 2019), applying one-sided Fisher's exact tests for each synaptic GO term, with FDR correction for multiple comparisons.

Gene expression profiles of $PLS^-$ and $PLS^+$ genes across human brain development were examined using whole-brain RNA-seq data from BrainSpan (http://www.brainspan.org/) (Hawrylycz et al., 2012). We obtained normalized gene expression values (RPKM) for 29 developmental time points from FUMA (https://fuma.ctglab.nl/) (Watanabe et al., 2017). These values were mean-centered per gene, such that positive values reflect expression above the gene's average across development, and negative values reflect expression below average. For each timepoint, we averaged expression values across PLS gene sets, and further summarized them across prenatal and postnatal stages to assess prenatal versus postnatal expression bias. Differences in expression between prenatal and postnatal stages were tested using Wilcoxon rank-sum tests.

To explore transcriptomic enrichment across tissues and cell types, we analyzed the overrepresentation of PLS genes among tissue- or cell-type-specific genes from two datasets: (1) GTEx v8, covering 13 brain regions and 29 non-brain tissues, and (2) single-cell RNA-sequencing data from 39 differentiated cell types across 19 mouse central and peripheral nervous system regions (GTEx Consortium, 2013; Zeisel et al., 2018). In both datasets, tissue- or cell-type-specific genes were defined as those in the top 10% of specificity values, following previous methods (Bryois et al., 2020). Overrepresentation of PLS genes was assessed by resampling, comparing observed overlaps to 10,000 simulated overlaps generated by random sampling from brain-expressed genes.

Genetic relationship between SZ-related sulcal width abnormalities and other related disorders

We leveraged data from differentially expressed genes (DEG) associated with psychiatric disorders from a prior study conducted by (Gandal et al., 2018). In this study, the authors used gene expression arrays' information, and applied a stringent normalization procedure and used technical covariates to address specific issues linked to working with post-mortem brain tissues (experimental batch, post-mortem interval, and pH). For this analysis, we considered genes as differentially expressed (DEG) when reaching nominal significance ($p < 0.05$) in each of the following disorders: schizophrenia (SCZ), bipolar disorder (BD), major depression (MDD), autism spectrum disorders (ASD), alcohol abuse disorder (ALC), and inflammatory bowel disease (IBD) as a non-psychiatric control. We assessed the overrepresentation of PLS+ and PLS- gene sets among DEG in each of these disorders. Overrepresentation analyses were performed across genes described to be up- or downregulated, separately. Enrichment significance was assessed by a resampling procedure, comparing real enrichment against enrichment distribution from 10,000 randomly background gene lists. *P*-values were corrected for multiple comparisons using FDR.

Enrichment for Common predisposing variation to SZ-related disorders and neuroimaging phenotypes

Enrichment of PLS- and PLS+ gene sets in common predisposing variation for SZ and other related disorders was assessed with MAGMA v1.10 (de Leeuw et al., 2015). First, enrichment of common variation was assessed across SZ and a wide range of related disorders/traits: Bipolar disorder (BD), Major Depression Disorder (MDD), Attention and deficit hyperactivity disorder (ADHD), Autism spectrum disorder (ASD), Obsessive-Compulsive Disorder (OCD), Alcohol Use Disorder (ALC) Cross-Disorder (CROSS), Neuroticism (NEUR), Cannabis Use Disorder (CUD), General Cognitive Function (COGN), Anorexia (ANO) and Tourette Syndrome (TS). Given the functional enrichments in neurodegenerative and energetic-synaptic functions, we also included in this list two disorders with well powered recent GWAS: Alzheimer (ALZ) and Parkinson disease (PARK). To control for non-specific polygenic signal, height GWAS (Yengo et al., 2022) was also included. Second, brain longitudinal phenotypes from ENIGMA Consortium GWAS (Brouwer and ENIGMA plasticity working group, 2022) with significant genetic overlap with SCZ were also studied (i.e, change in CT, total brain volume, hippocampus, surface area, nucleus accumbens, and cerebellum white matter). Additionally, we also included Functional and Structural Connectivity phenotypes within cerebral resting-state networks from a recent GWAS (Tissink et al., 2023). Enrichment was evaluated by one-tailed competitive test, using the list of 15,633 brain-expressed genes as background and gene boundaries for variant inclusion (35 kb upstream and 10 kb downstream). GWAS summary data used is described in Supplemental Data 2.

Common and independent biological functionalities of SZ-related sulcal width and cortical thickness abnormalities

We investigated whether the biological functions of genes whose anatomically informed expression patterns were associated with sulcal widening in SZ overlap with—or are distinct from—those linked to baseline cortical thickness (CT) differences or to accelerated cortical thinning (ACT), as previously described in SZ. To this end, we compared the PLS gene sets associated with sulcal widening in the present study ($PLS^-$, $PLS^+$) with those associated with CT differences ($CTD\text{-}PLS^-$, $CTD\text{-}PLS^+$) and ACT ($ACT\text{-}PLS^-$, $ACT\text{-}PLS^+$) as identified in our prior work (González-Peñas et al., 2024). Overlap between gene sets was evaluated using 10,000 random permutations drawn from the common background of brain-expressed genes, with odds ratios (ORs) computed via Fisher's exact test and *p*-values corrected for multiple comparisons using FDR.

Among the comparisons, significant overlap was observed only between the $ACT\text{-}PLS^-$ and $PLS^-$ gene sets. These two sets also showed the strongest functional, transcriptional, and genetic enrichments among all PLS-defined groups. However, the actual overlap was modest (66 of 1,001 $PLS^-$ genes), prompting us to assess whether the observed enrichments were driven by this subset. We therefore repeated all major enrichment analyses on the remaining 934 $PLS^-$ genes that did not overlap with $ACT\text{-}PLS^-$ genes.

Despite the reduction in gene count (with 13,146 genes in common across studies), the non-overlapping $PLS^-$ genes showed similar significant enrichment for mitochondrial and energy metabolism pathways, including aerobic respiration ($p_{FDR}$ =

1.95 × $10^{-5}$), ATP synthesis ($p_{FDR}$ = 0.02), mitochondrial transport ($p_{FDR}$ = 5.42 × $10^{-3}$), and glycolysis/gluconeogenesis ($p_{FDR}$ = 0.016) (see Supplementary Data 12). Disease-association analyses linked these genes to metabolic dysfunctions such as elevated serum lactate ($p_{FDR}$ = 0.002) and lactic acidosis ($p_{FDR}$ = 0.007). As with the full $PLS^-$ set, the non-overlapping $PLS^-$ genes were also enriched for synaptic functions—particularly at the presynapse ($p_{FDR}$ = 0.034), in synaptic vesicle cycling ($p_{FDR}$ = 0.019), and synaptic transport ($p_{FDR}$ = 0.02), as defined by SynGO annotations (see Supplementary Data 13).

Transcriptional profiling of the non-overlapping $PLS^-$ genes revealed enrichment patterns nearly identical to those of the full set: genes were highly expressed in cortical and hypothalamic tissue, and, beyond the brain, in cardiac and muscle tissues (see Supplementary Data 14). At the cellular level, enrichments were found in peripheral sensory neurons (both peptidergic and non-peptidergic), cholinergic and monoaminergic neurons, and hindbrain neurons involved in sensory-motor integration (see Supplementary Data 15). Additionally, these non-overlapping $PLS^-$ genes were significantly enriched among genes downregulated in SZ, bipolar disorder (BD), and autism spectrum disorder (ASD), replicating findings based on the complete $PLS^-$ set (see Supplementary Data 16).

Together, these results demonstrate that the transcriptional architecture associated with sulcal widening in SZ is largely independent from that related to baseline CT or ACT. The genes associated with sulcal widening display unique functional and spatial characteristics—marked by synaptic and metabolic roles, neuronal specificity, and cortico-hypothalamic expression profiles—underscoring the distinct biological processes underlying this anatomical phenotype.

Supplementary Table 2. Balanced classification accuracy scores (%) from the linear support vector machine predicting samples within healthy controls from the train and test cohorts.

| Site | Train Samples | Test Samples | Train | Test |
|---|---|---|---|---|
| camcan | 258 | 258 | 0.48 | 0.45 |
| oasis | 539 | 539 | 0.48 | 0.47 |
| ixi | 502 | 502 | 0.47 | 0.45 |
| aomic_piop2 | 218 | 218 | 0.45 | 0.45 |
| dlbs | 145 | 145 | 0.45 | 0.41 |
| narratives | 325 | 325 | 0.46 | 0.49 |
| sald | 450 | 450 | 0.47 | 0.46 |
| aomic_piop1 | 196 | 196 | 0.43 | 0.42 |
| rockland | 792 | 792 | 0.47 | 0.46 |
| aomic_id1000 | 886 | 886 | 0.47 | 0.48 |
| cobre | 122 | 122 | 0.49 | 0.52 |
| bgs | 120 | 120 | 0.52 | 0.51 |

| | | | | |
|---|---|---|---|---|
| utrecht | 386 | 386 | 0.52 | 0.52 |
| SRPBS_open | 300 | 300 | 0.50 | 0.51 |
| LA5C_study | 153 | 153 | 0.49 | 0.46 |

Supplementary Table 3. For each sulcus, group means, t-values, uncorrected p-values and Cohen's d-values for the diagnostic group difference of normative modeling-based z-scores of sulcal width. P-values significant after FDR correction for multiple comparisons are given in bold.

| Sulcus | mean $HC_{test}$ | mean SZ | *t* | *p* | *d* |
|---|---|---|---|---|---|
| Left Central Sulcus | 0.07 | 0.17 | -1.46 | 0.15 | -0.10 |
| Left Pre-Central Sulcus | 0.03 | 0.43 | -6.12 | **<0.001** | -0.41 |
| Left Post-Central Sulcus | 0.18 | 0.17 | 0.11 | 0.91 | 0.01 |
| Left Intra-Parietal Fissure | 0.01 | 0.04 | -0.39 | 0.69 | -0.03 |
| Left Superior Frontal Sulcus | -0.01 | 0.33 | -5.10 | **<0.001** | -0.34 |
| Left Parieto-Occipital Fissure | 0.01 | 0.08 | -1.07 | 0.28 | -0.07 |
| Left Posterior Cingulate Sulcus | 0.01 | -0.03 | 0.51 | 0.61 | 0.03 |
| Left Paracingulate Sulcus | 0.04 | 0.22 | -2.67 | **0.01** | -0.18 |
| Left Inferior Frontal Sulcus | 0.00 | 0.50 | -7.74 | **<0.001** | -0.52 |
| Left Orbitofrontal Sulcus | 0.05 | 0.31 | -3.66 | **<0.001** | -0.25 |
| Left Lateral Fissure | -0.09 | 0.49 | -8.28 | **<0.001** | -0.55 |
| Left Anterior Cingulate Sulcus | 0.04 | 0.22 | -2.55 | **0.01** | -0.17 |
| Left Superior Temporal Sulcus | 0.08 | 0.48 | -6.14 | **<0.001** | -0.41 |
| Left Occipital Sulcus | -0.02 | 0.24 | -3.61 | **<0.001** | -0.25 |
| Left Collateral Fissure | 0.02 | 0.47 | -7.01 | **<0.001** | -0.46 |
| Left Medial Parietal Sulcus | 0.04 | 0.06 | -0.33 | 0.74 | -0.02 |
| Left Intermediate Frontal Sulcus | 0.04 | 0.41 | -5.15 | **<0.001** | -0.34 |
| Left Inferior Temporal Sulcus | -0.07 | 0.49 | -8.23 | **<0.001** | -0.55 |
| Left Occipital-Temporal Lateral Sulcus | 0.02 | 0.42 | -5.66 | **<0.001** | -0.39 |
| Left Calcarine Fissure | 0.08 | 0.44 | -4.68 | **<0.001** | -0.32 |
| Right Central Sulcus | 0.03 | 0.09 | -0.75 | 0.45 | -0.05 |
| Right Pre-Central Sulcus | 0.00 | 0.39 | -5.73 | **<0.001** | -0.38 |
| Right Post-Central Sulcus | 0.08 | 0.07 | 0.14 | 0.89 | 0.01 |
| Right Intra-Parietal Fissure | 0.08 | 0.11 | -0.44 | 0.66 | -0.03 |
| Right Superior Frontal Sulcus | 0.06 | 0.33 | -4.17 | **<0.001** | -0.28 |
| Right Parieto-Occipital Fissure | -0.06 | 0.14 | -2.96 | **0.00** | -0.20 |
| Right Posterior Cingulate Sulcus | -0.02 | -0.02 | -0.13 | 0.90 | -0.01 |
| Right Paracingulate Sulcus | -0.00 | 0.34 | -4.93 | **<0.001** | -0.33 |
| Right Inferior Frontal Sulcus | 0.01 | 0.71 | -10.25 | **<0.001** | -0.69 |
| Right Orbitofrontal Sulcus | 0.03 | 0.26 | -3.32 | **0.01** | -0.22 |
| Right Lateral Fissure | -0.05 | 0.47 | -7.95 | **<0.001** | -0.54 |

| | | | | | |
|---|---|---|---|---|---|
| Right Anterior Cingulate Sulcus | 0.08 | 0.28 | -2.92 | **0.01** | -0.20 |
| Right Superior Temporal Sulcus | 0.08 | 0.39 | -4.69 | **<0.001** | -0.32 |
| Right Occipital Sulcus | -0.02 | 0.29 | -4.32 | **<0.001** | -0.29 |
| Right Collateral Fissure | 0.05 | 0.48 | -6.37 | **<0.001** | -0.43 |
| Right Medial Parietal Sulcus | 0.09 | 0.07 | 0.18 | 0.85 | 0.01 |
| Right Intermediate Frontal Sulcus | -0.01 | 0.44 | -6.73 | **<0.001** | -0.45 |
| Right Inferior Temporal Sulcus | -0.10 | 0.43 | -7.25 | **<0.001** | -0.49 |
| Right Occipital-Temporal Lateral Sulcus | -0.12 | 0.25 | -5.40 | **<0.001** | -0.36 |
| Right Calcarine Fissure | -0.06 | 0.33 | -5.48 | **<0.001** | -0.37 |

Supplementary Table 4. For each sulcus, group percentages, **$X^2$**-values from permuted chi-square tests (n=10,000), uncorrected p-values and Cohen's ω-values (similar interpretation as Cohen's *d* values) for the diagnostic group difference of the proportion of individuals with infra-normal (extreme negative deviance) values. P-values significant after FDR correction for multiple comparisons are given in bold.

| Sulcus | Proportion Infra $HC_{test}$ | Proportion Infra SZ | $X^2$ | *p* | ω |
|---|---|---|---|---|---|
| Left Central Sulcus | 0.02 | 0.01 | 1.10 | 0.29 | 0.03 |
| Left Pre-Central Sulcus | 0.02 | 0.02 | 0.08 | 0.77 | 0.01 |
| Left Post-Central Sulcus | 0.02 | 0.03 | 0.60 | 0.44 | 0.03 |
| Left Intra-Parietal Fissure | 0.01 | 0.02 | 0.73 | 0.39 | 0.03 |
| Left Superior Frontal Sulcus | 0.03 | 0.02 | 1.05 | 0.31 | 0.03 |
| Left Parieto-Occipital Fissure | 0.03 | 0.03 | 0.01 | 0.93 | 0.00 |
| Left Posterior Cingulate Sulcus | 0.03 | 0.04 | 0.58 | 0.45 | 0.03 |
| Left Paracingulate Sulcus | 0.02 | 0.03 | 0.61 | 0.43 | 0.03 |
| Left Inferior Frontal Sulcus | 0.03 | 0.02 | 0.92 | 0.34 | 0.03 |
| Left Orbitofrontal Sulcus | 0.02 | 0.02 | 0.42 | 0.51 | 0.02 |
| Left Lateral Fissure | 0.03 | 0.01 | 1.97 | 0.16 | 0.05 |
| Left Anterior Cingulate Sulcus | 0.03 | 0.03 | 0.26 | 0.61 | 0.02 |
| Left Superior Temporal Sulcus | 0.03 | 0.03 | 0.04 | 0.84 | 0.01 |
| Left Occipital Sulcus | 0.04 | 0.04 | 0.01 | 0.93 | 0.00 |
| Left Collateral Fissure | 0.02 | 0.01 | 2.55 | 0.11 | 0.05 |
| Left Medial Parietal Sulcus | 0.03 | 0.03 | 0.00 | 0.99 | 0.00 |
| Left Intermediate Frontal Sulcus | 0.05 | 0.01 | 9.43 | 0.00 | 0.10 |
| Left Inferior Temporal Sulcus | 0.04 | 0.01 | 4.13 | 0.04 | 0.07 |
| Left Occipital-Temporal Lateral Sulcus | 0.03 | 0.01 | 1.58 | 0.21 | 0.04 |
| Left Calcarine Fissure | 0.02 | 0.02 | 0.00 | 1.00 | 0.00 |
| Right Central Sulcus | 0.04 | 0.03 | 1.43 | 0.23 | 0.04 |
| Right Pre-Central Sulcus | 0.03 | 0.01 | 1.98 | 0.16 | 0.05 |
| Right Post-Central Sulcus | 0.03 | 0.02 | 0.13 | 0.72 | 0.01 |
| Right Intra-Parietal Fissure | 0.02 | 0.03 | 0.86 | 0.35 | 0.03 |
| Right Superior Frontal Sulcus | 0.02 | 0.02 | 0.46 | 0.50 | 0.02 |
| Right Parieto-Occipital Fissure | 0.02 | 0.03 | 0.43 | 0.51 | 0.02 |
| Right Posterior Cingulate Sulcus | 0.02 | 0.04 | 3.57 | 0.06 | 0.06 |
| Right Paracingulate Sulcus | 0.02 | 0.01 | 1.32 | 0.25 | 0.04 |
| Right Inferior Frontal Sulcus | 0.02 | 0 | 5.11 | 0.02 | 0.08 |

| Sulcus | Proportion Infra $HC_{test}$ | Proportion Infra SZ | $\chi^2$ | *p* | ω |
|---|---|---|---|---|---|
| Right Orbitofrontal Sulcus | 0.02 | 0.02 | 0.16 | 0.69 | 0.01 |
| Right Lateral Fissure | 0.03 | 0.01 | 3.68 | 0.05 | 0.06 |
| Right Anterior Cingulate Sulcus | 0.01 | 0.01 | 0.08 | 0.77 | 0.01 |
| Right Superior Temporal Sulcus | 0.03 | 0.02 | 0.72 | 0.40 | 0.03 |
| Right Occipital Sulcus | 0.05 | 0.02 | 3.24 | 0.07 | 0.06 |
| Right Collateral Fissure | 0.02 | 0.02 | 0.00 | 0.94 | 0.00 |
| Right Medial Parietal Sulcus | 0.03 | 0.03 | 0.01 | 0.91 | 0.00 |
| Right Intermediate Frontal Sulcus | 0.04 | 0.01 | 5.68 | 0.02 | 0.08 |
| Right Inferior Temporal Sulcus | 0.04 | 0.01 | 5.72 | 0.02 | 0.08 |
| Right Occipital-Temporal Lateral Sulcus | 0.05 | 0.01 | 7.21 | 0.01 | 0.09 |
| Right Calcarine Fissure | 0.02 | 0.02 | 0.10 | 0.75 | 0.01 |

Supplementary Table 5. For each sulcus, group percentages, **$X^2$**-values from permuted chi-square tests (n=10,000), uncorrected p-values and Cohen's ω-values for the diagnostic group difference of the proportion of individuals with supra-normal (extreme positive deviance) values. P-values significant after FDR correction for multiple comparisons are given in bold.

| Sulcus | Proportion Supra $HC_{test}$ | Proportion Supra SZ | $X^2$ | *p* | ω |
|---|---|---|---|---|---|
| Left Central Sulcus | 0.04 | 0.05 | 0.00 | 0.95 | 0.00 |
| Left Pre-Central Sulcus | 0.02 | 0.06 | 14.14 | **<0.001** | 0.12 |
| Left Post-Central Sulcus | 0.04 | 0.04 | 0.48 | 0.49 | 0.02 |
| Left Intra-Parietal Fissure | 0.03 | 0.03 | 0.27 | 0.60 | 0.02 |
| Left Superior Frontal Sulcus | 0.02 | 0.04 | 3.40 | 0.07 | 0.06 |
| Left Parieto-Occipital Fissure | 0.03 | 0.02 | 0.71 | 0.40 | 0.03 |
| Left Posterior Cingulate Sulcus | 0.05 | 0.04 | 0.16 | 0.69 | 0.01 |
| Left Paracingulate Sulcus | 0.04 | 0.06 | 1.60 | 0.21 | 0.04 |
| Left Inferior Frontal Sulcus | 0.02 | 0.07 | 16.44 | **<0.001** | 0.13 |
| Left Orbitofrontal Sulcus | 0.03 | 0.07 | 8.81 | **0.00** | 0.10 |
| Left Lateral Fissure | 0.03 | 0.1 | 17.90 | **<0.001** | 0.14 |
| Left Anterior Cingulate Sulcus | 0.04 | 0.04 | 0.10 | 0.75 | 0.01 |
| Left Superior Temporal Sulcus | 0.02 | 0.06 | 10.38 | **0.00** | 0.11 |
| Left Occipital Sulcus | 0.02 | 0.06 | 8.02 | **0.00** | 0.09 |
| Left Collateral Fissure | 0.03 | 0.07 | 5.37 | **0.02** | 0.08 |
| Left Medial Parietal Sulcus | 0.03 | 0.03 | 0.02 | 0.90 | 0.00 |
| Left Intermediate Frontal Sulcus | 0.04 | 0.06 | 1.23 | 0.27 | 0.04 |
| Left Inferior Temporal Sulcus | 0.02 | 0.08 | 17.35 | **<0.001** | 0.14 |
| Left Occipital-Temporal Lateral Sulcus | 0.02 | 0.07 | 14.35 | **<0.001** | 0.12 |
| Left Calcarine Fissure | 0.04 | 0.1 | 17.01 | **<0.001** | 0.14 |
| Right Central Sulcus | 0.04 | 0.05 | 1.68 | 0.19 | 0.04 |
| Right Pre-Central Sulcus | 0.02 | 0.07 | 12.99 | **0.00** | 0.12 |
| Right Post-Central Sulcus | 0.04 | 0.03 | 1.03 | 0.31 | 0.03 |
| Right Intra-Parietal Fissure | 0.04 | 0.03 | 0.27 | 0.60 | 0.02 |
| Right Superior Frontal Sulcus | 0.02 | 0.05 | 5.20 | **0.02** | 0.07 |
| Right Parieto-Occipital Fissure | 0.02 | 0.04 | 2.87 | 0.09 | 0.06 |
| Right Posterior Cingulate Sulcus | 0.02 | 0.03 | 0.52 | 0.47 | 0.02 |
| Right Paracingulate Sulcus | 0.03 | 0.07 | 6.96 | **0.01** | 0.09 |
| Right Inferior Frontal Sulcus | 0.03 | 0.1 | 22.83 | **<0.001** | 0.16 |

| | | | | | |
|---|---|---|---|---|---|
| Right Orbitofrontal Sulcus | 0.03 | 0.05 | 2.10 | 0.15 | 0.05 |
| Right Lateral Fissure | 0.02 | 0.08 | 15.16 | **<0.001** | 0.13 |
| Right Anterior Cingulate Sulcus | 0.05 | 0.05 | 0.01 | 0.92 | 0.00 |
| Right Superior Temporal Sulcus | 0.03 | 0.03 | 0.03 | 0.87 | 0.01 |
| Right Occipital Sulcus | 0.04 | 0.06 | 2.18 | 0.14 | 0.05 |
| Right Collateral Fissure | 0.03 | 0.09 | 16.46 | **<0.001** | 0.13 |
| Right Medial Parietal Sulcus | 0.03 | 0.04 | 0.16 | 0.69 | 0.01 |
| Right Intermediate Frontal Sulcus | 0.02 | 0.06 | 7.58 | **0.01** | 0.09 |
| Right Inferior Temporal Sulcus | 0.02 | 0.07 | 15.33 | **<0.001** | 0.13 |
| Right Occipital-Temporal Lateral Sulcus | 0.03 | 0.05 | 2.76 | 0.10 | 0.05 |
| Right Calcarine Fissure | 0.02 | 0.08 | 16.76 | **<0.001** | 0.13 |

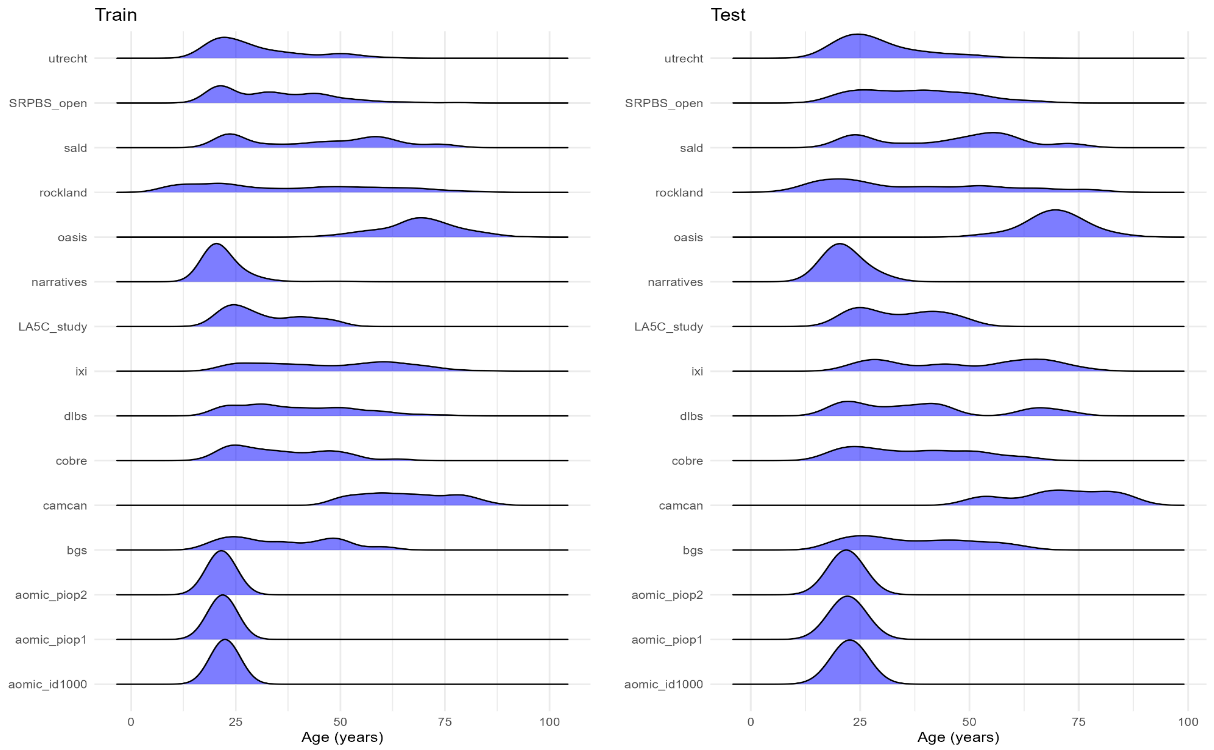


Supplementary Figure 1. The age distribution for each dataset within the training/test set. For details on each dataset see Supplementary Table 1 (part of the Supplementary Data).

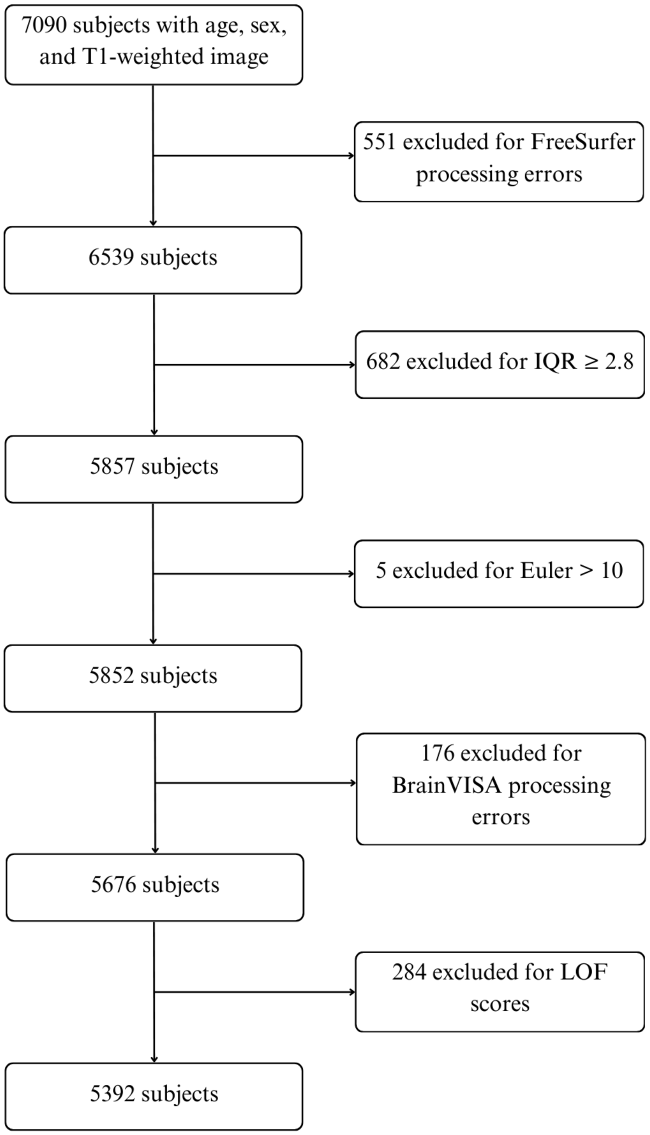


Supplementary Figure 2. Flow diagram showing the inclusion/exclusion process.

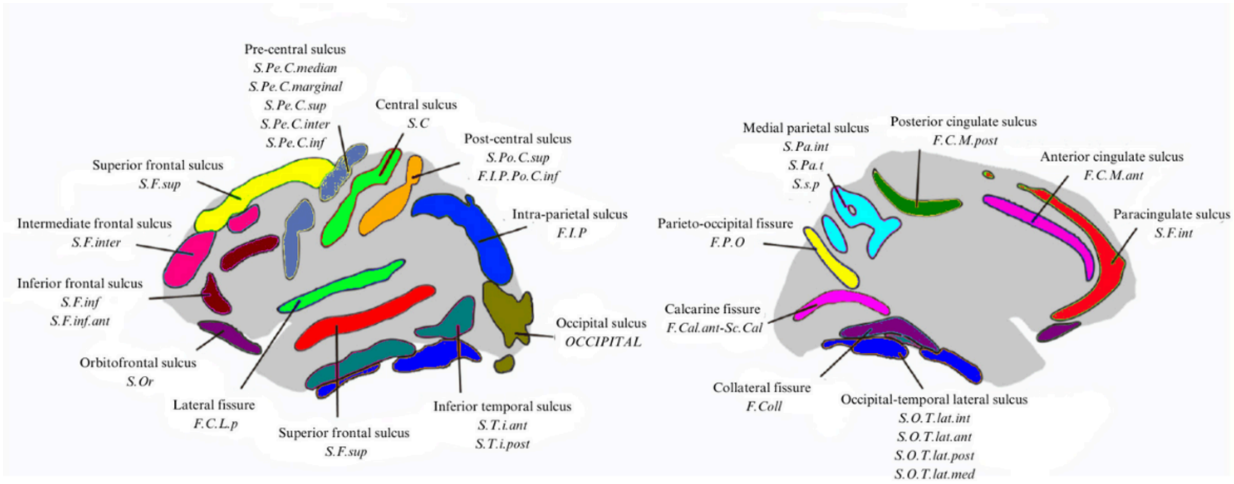


Supplementary Figure 3. Sulcal nomenclature was adapted from the BrainVISA Morphologist framework. Lateral and medial cortical views are shown, with sulci in colors. The sulcal labels used in this study follow the taxonomy proposed by Snyder et al. (2024) (Snyder et al., 2024) and the standard BrainVISA Morphologist taxonomy, with French abbreviations italicized beneath each label.

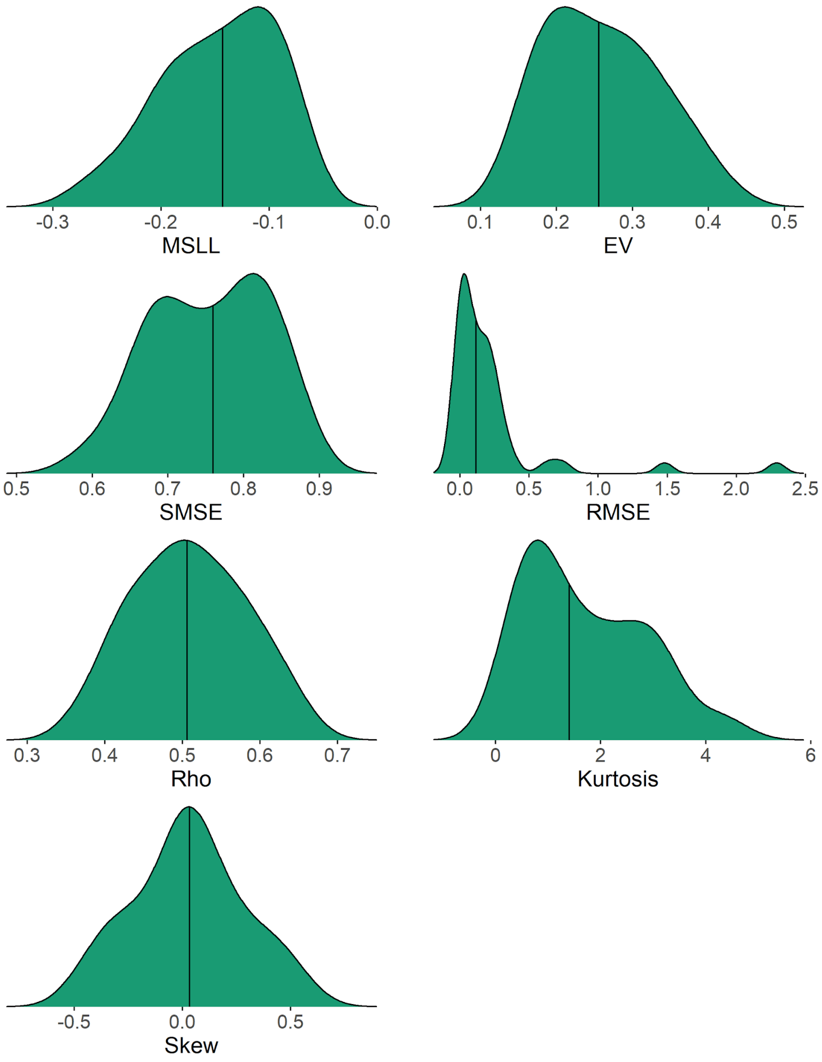


Supplementary Figure 4. Performance metrics of normative models. The distribution and the median for the performance metrics for the normative models, assessing their accuracy in estimating the relationship between sulcal width and age, sex, Euler number and scanner. These metrics encompass i) Mean standardized log-loss (MSLL, lower values indicating better performance), ii) explained variance (EV, higher values indicating better performance), iii) Standardized mean squared error (SMSE, lower values indicating better performance), iv) Root mean squared error (RMSE, lower values indicating better performance), v) Rho (higher values indicate better performance), vi) Kurtosis and vii) Skewness (Skew). Q-Q plots for all normative models are available from https://github.com/iamjoostjanssen/NormModel_Sulci/tree/main/Q-Q%

Supplementary Figure 5. Centile plots from normative modeling of sulcal width for each sulcus (continues on next page).

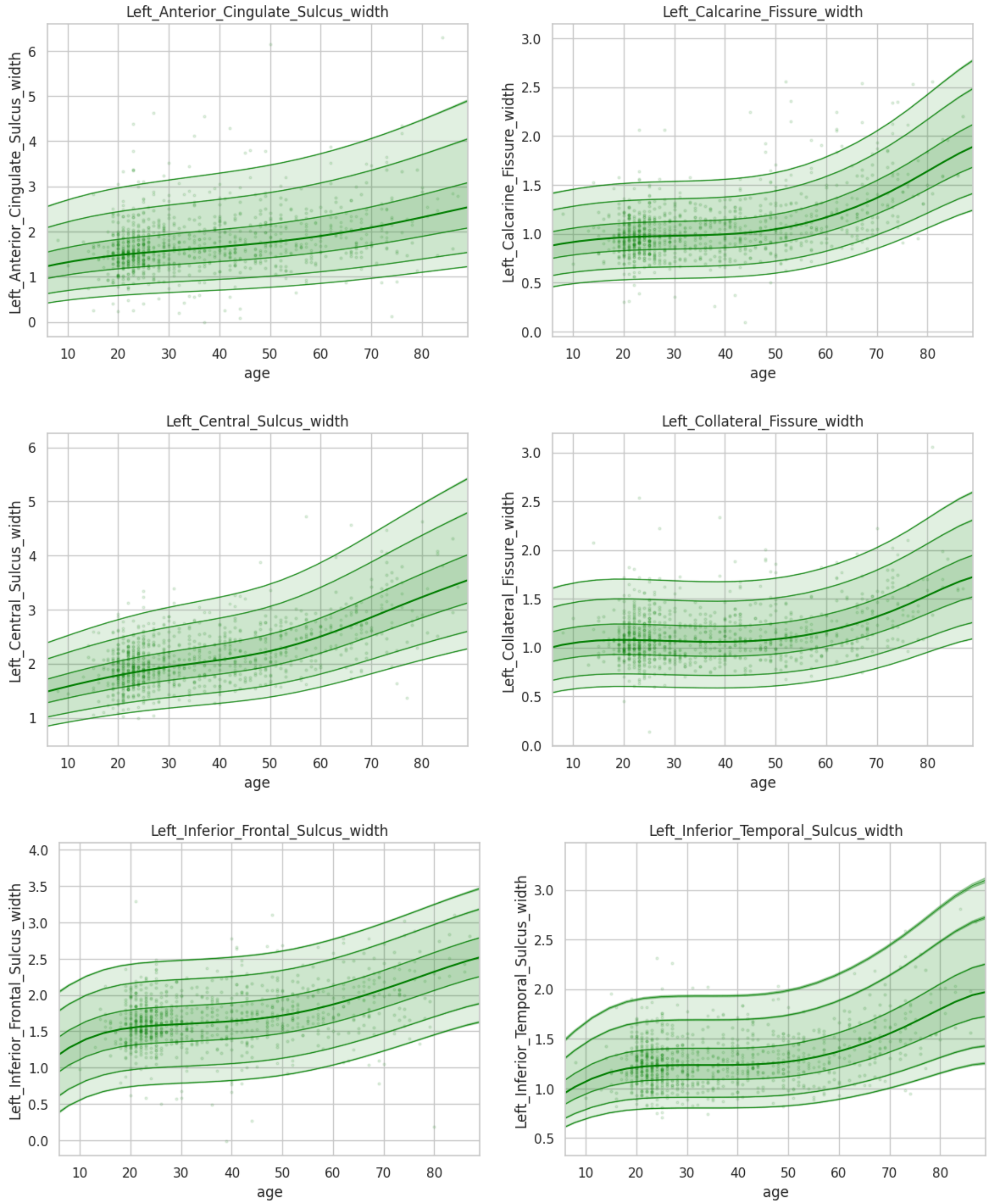

Supplementary Figure 5. Continued.

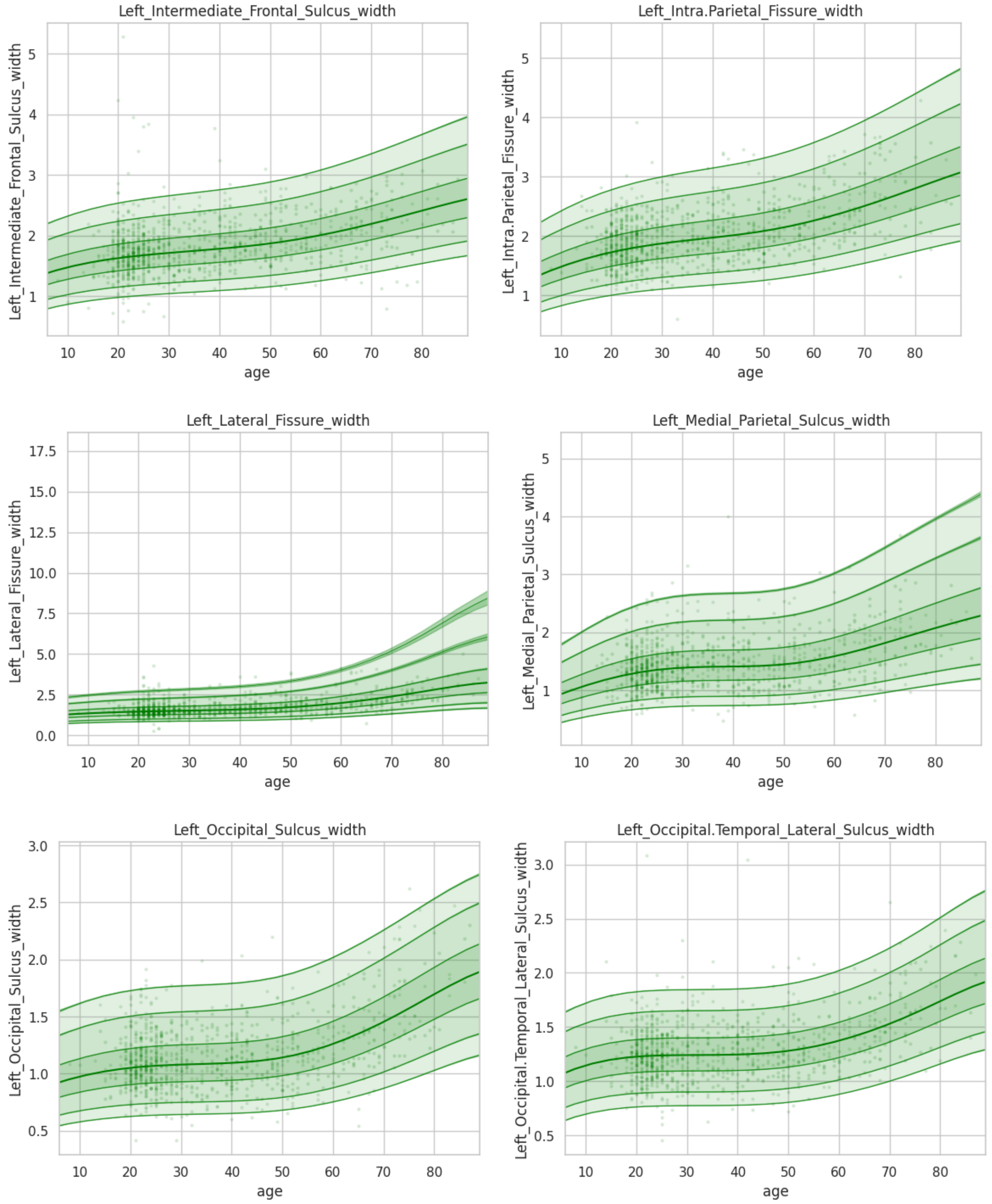

Supplementary Figure 5. Continued.

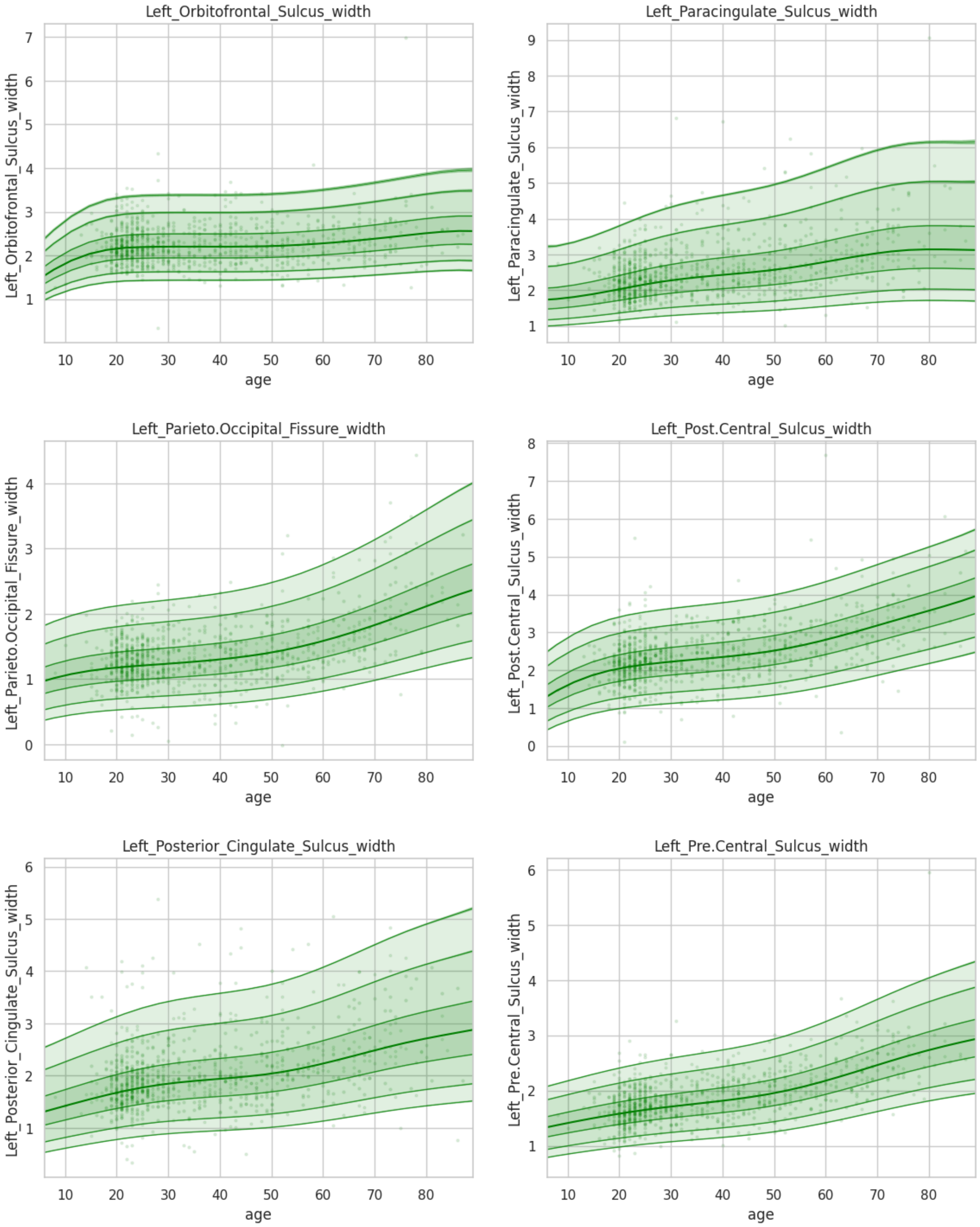

Supplementary Figure 5. Continued.

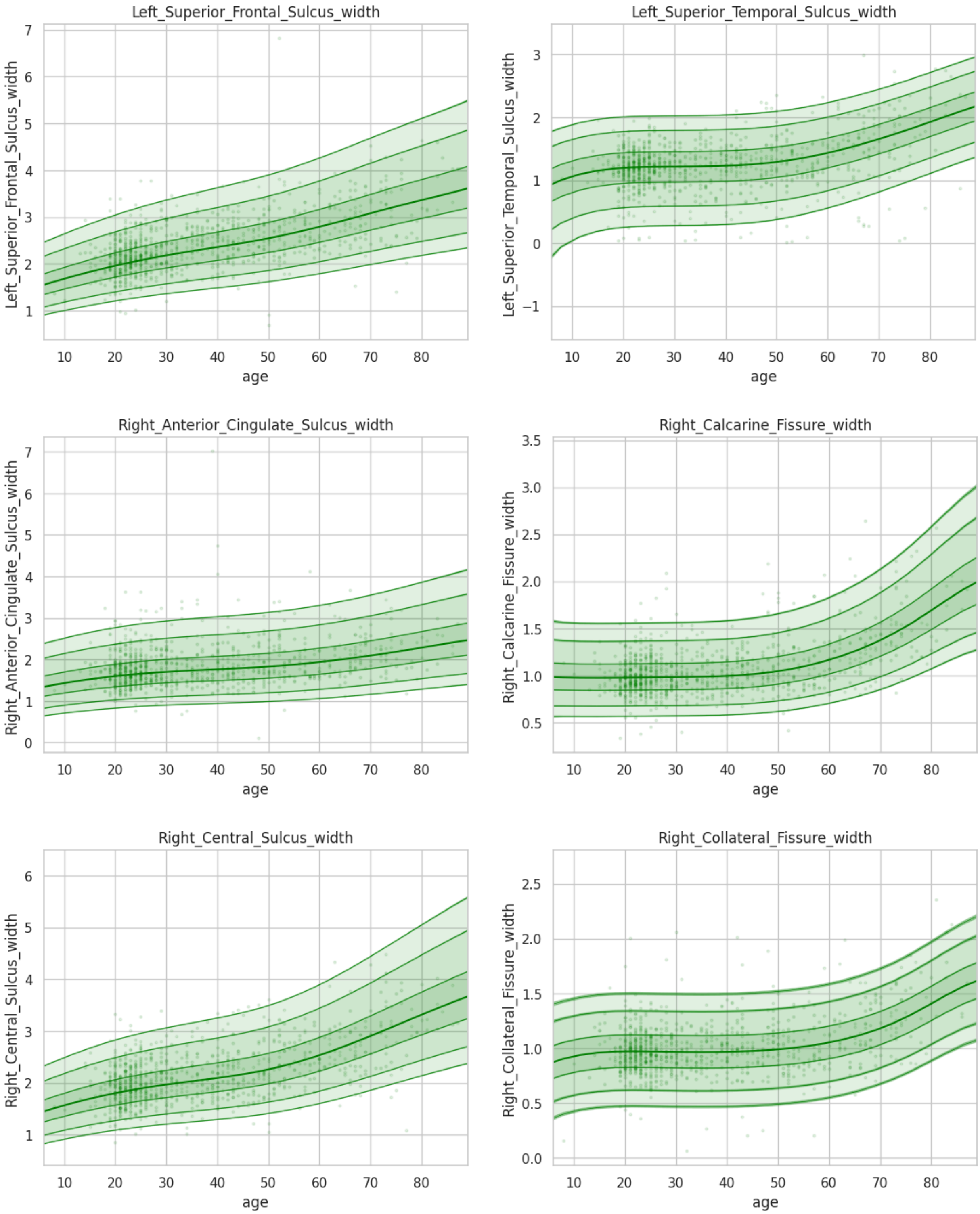

Supplementary Figure 5. Continued.

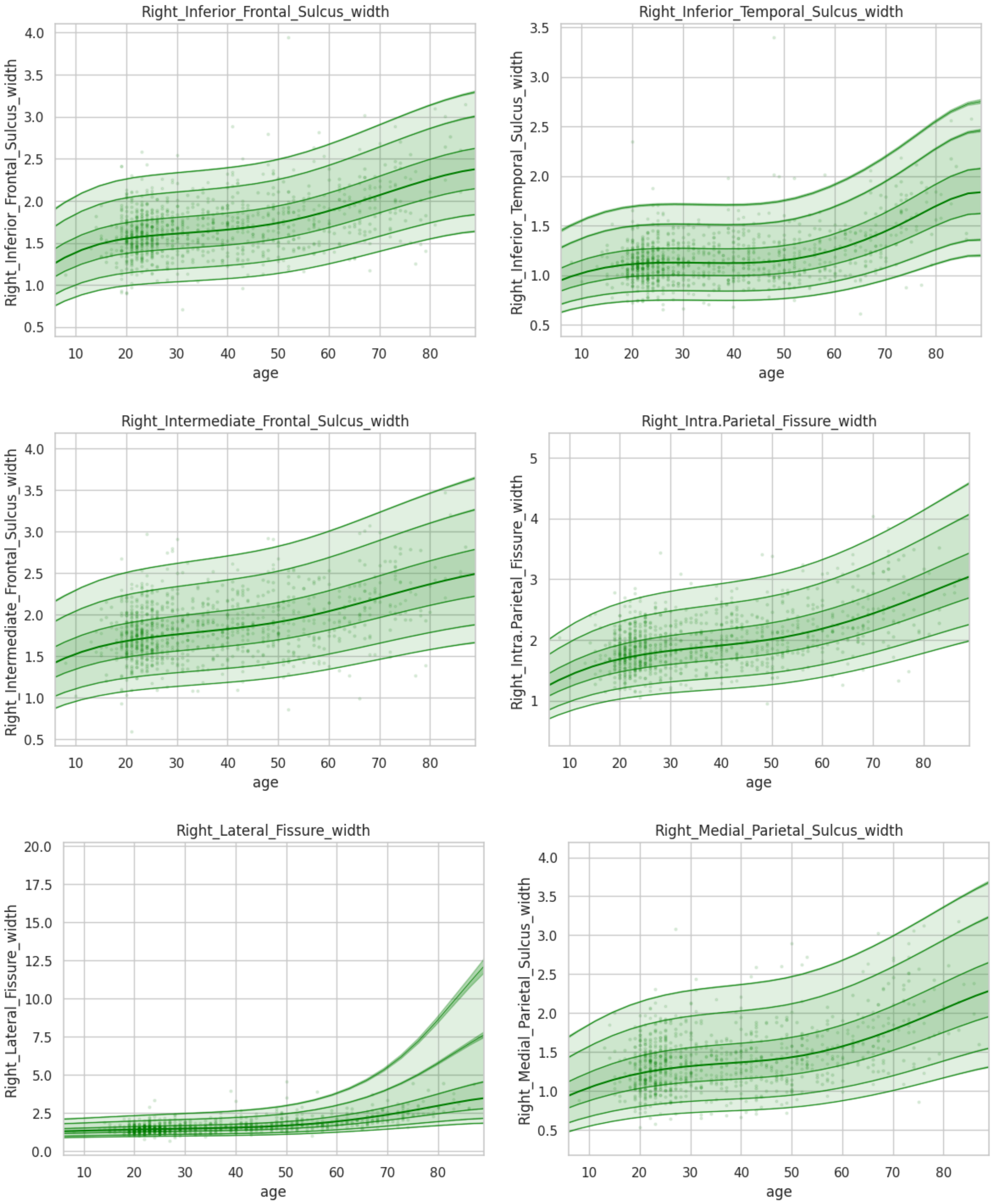

Supplementary Figure 5. Continued.

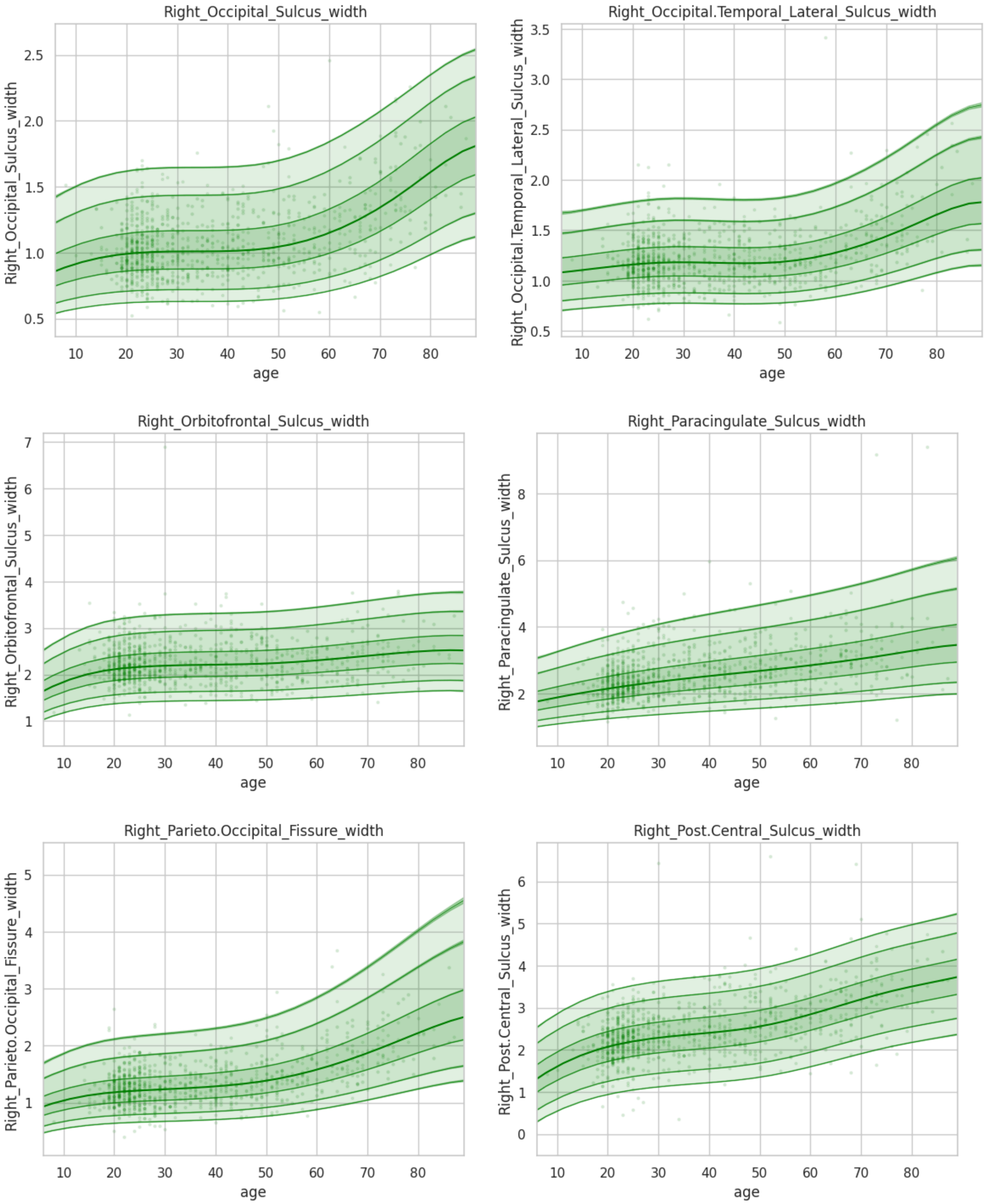

Supplementary Figure 5. Continued.

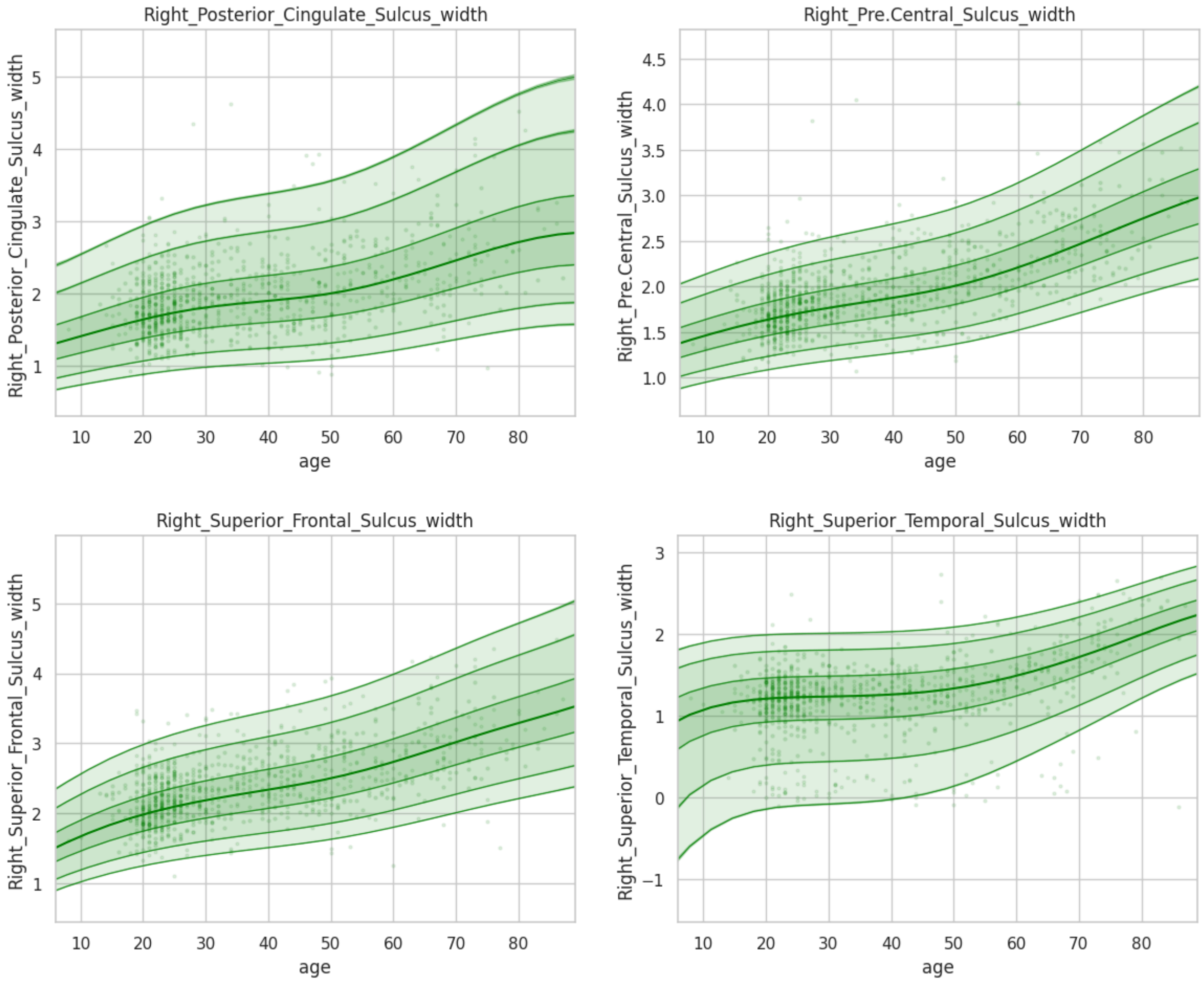

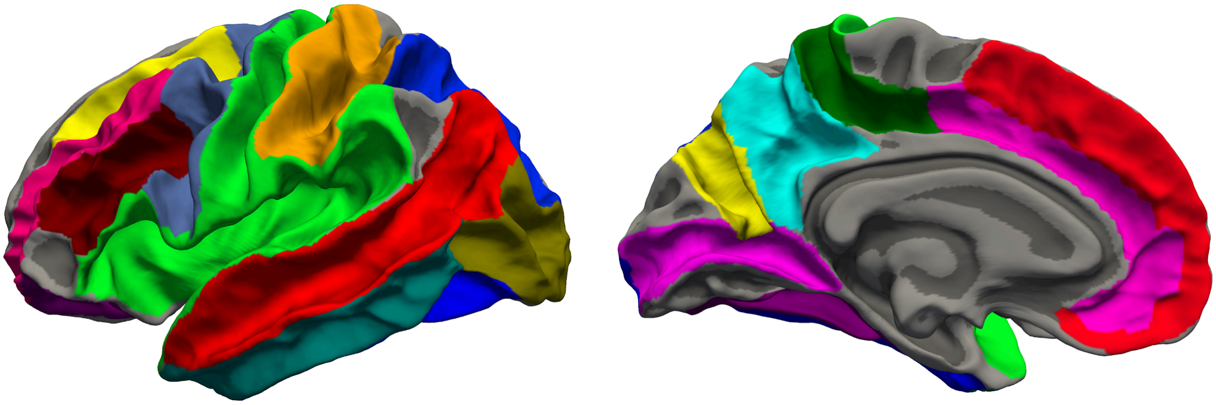

Supplementary Figure 6. Sulcal parcellation. The left hemispheric sulcal parcellation in 'fsaverage' space was designed so that each sulcal basin was centered on the sulcal fundus and extended up through the sulcal banks, with boundaries between regions at gyral ridges.

Supplementary Figure 7. Individual z-scores and diagnostic group averages (continues on next page).

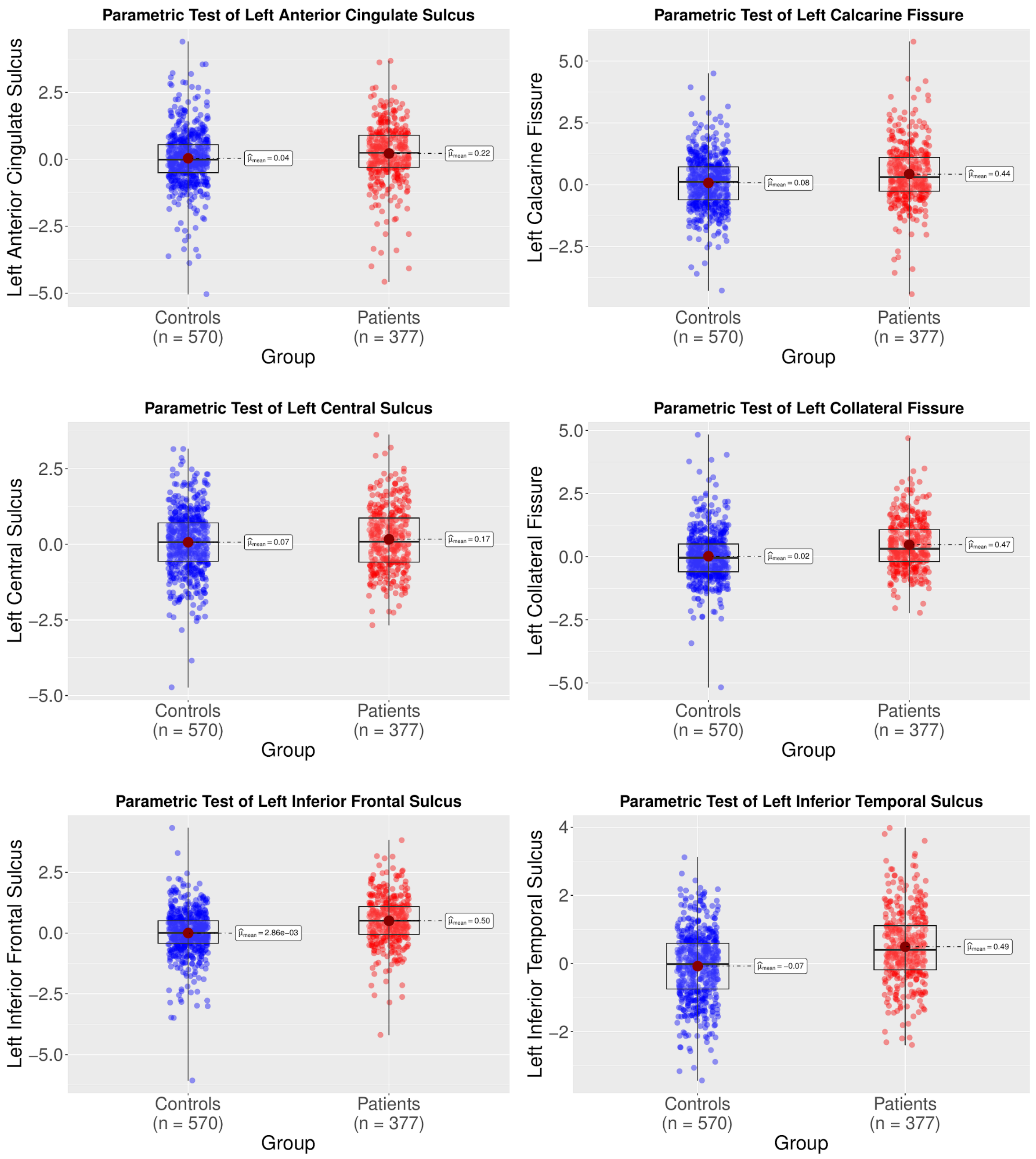

Supplementary Figure 7. Continued.

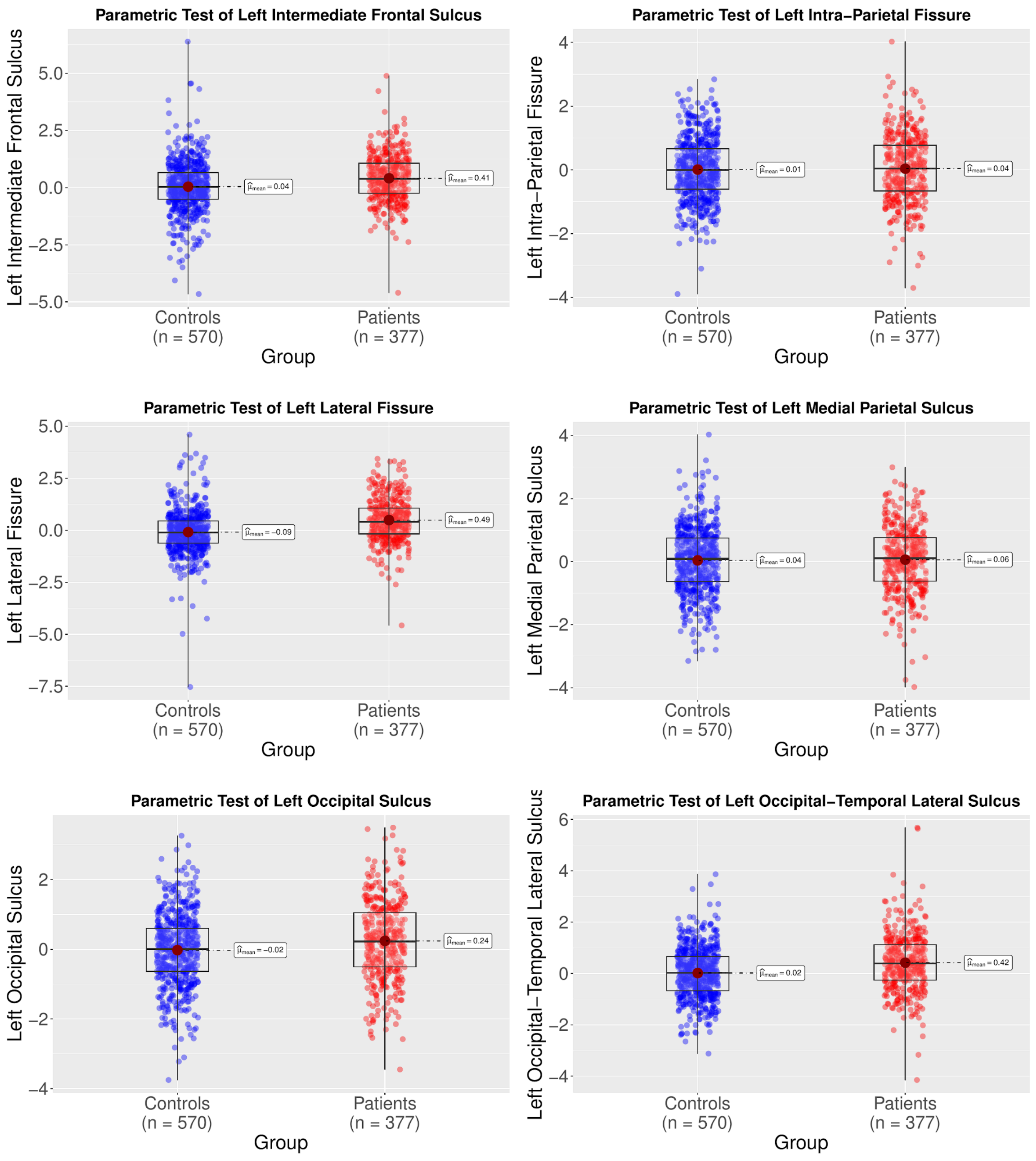

Supplementary Figure 7. Continued.

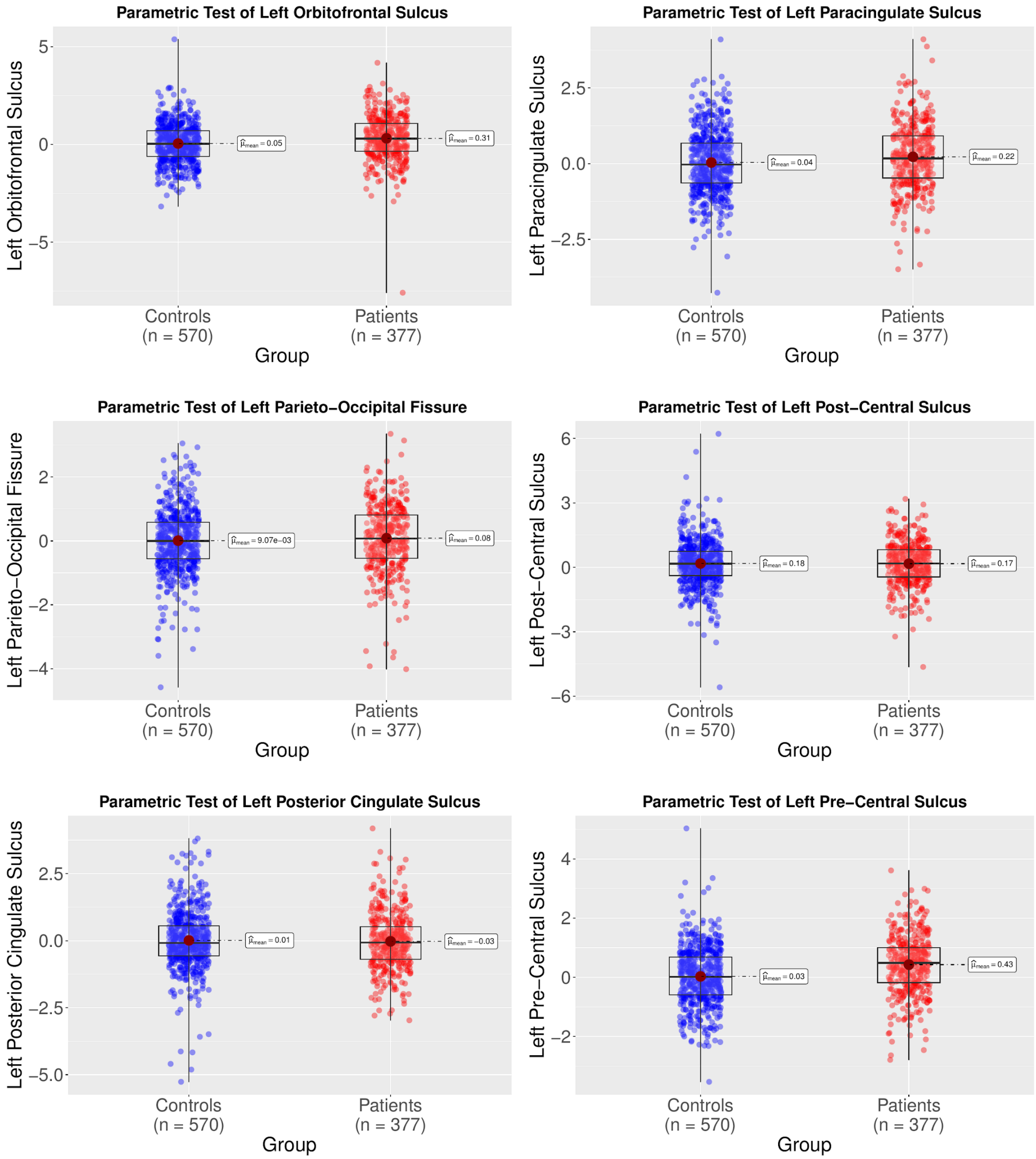

Supplementary Figure 7. Continued.

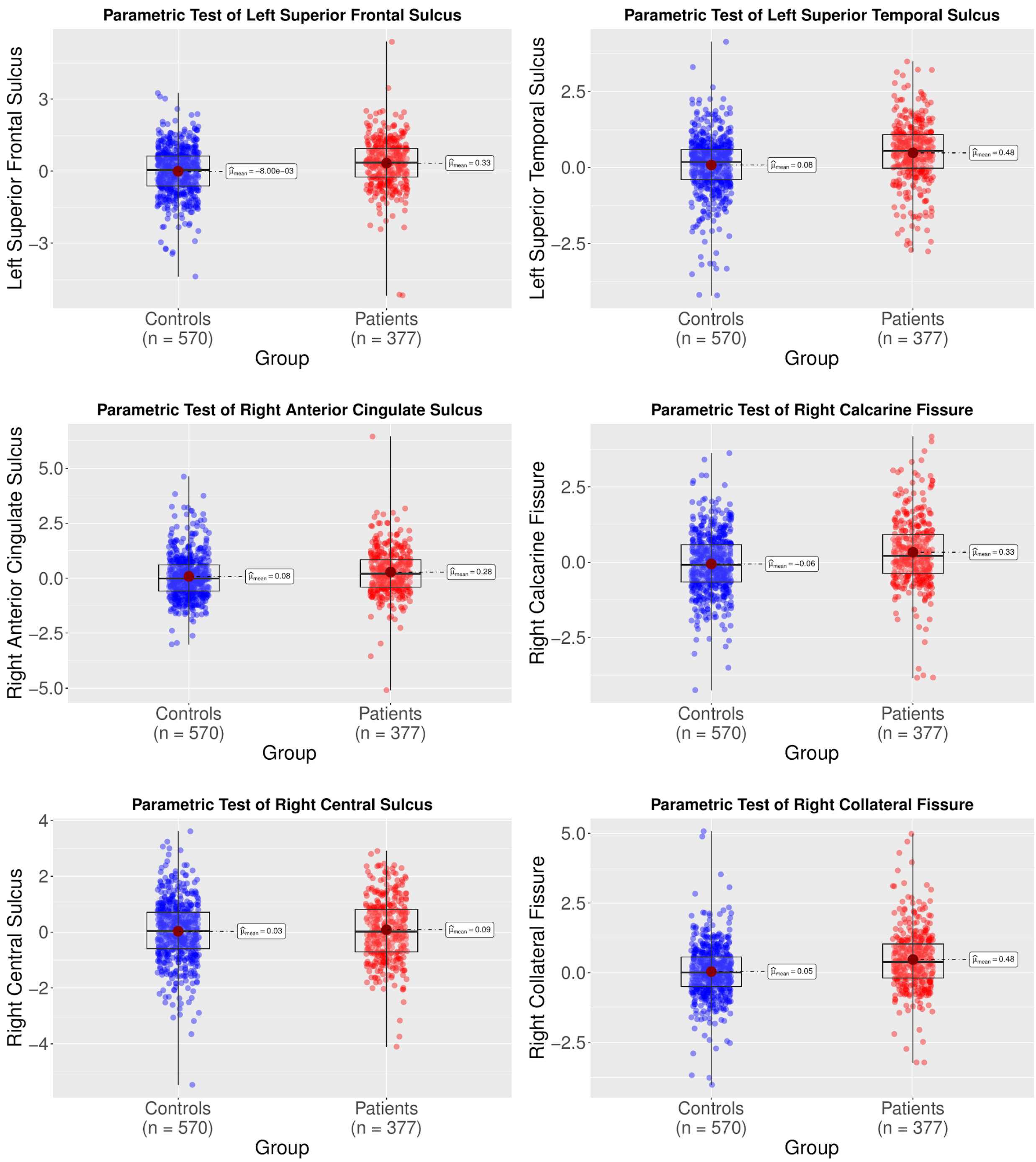

Supplementary Figure 7. Continued.

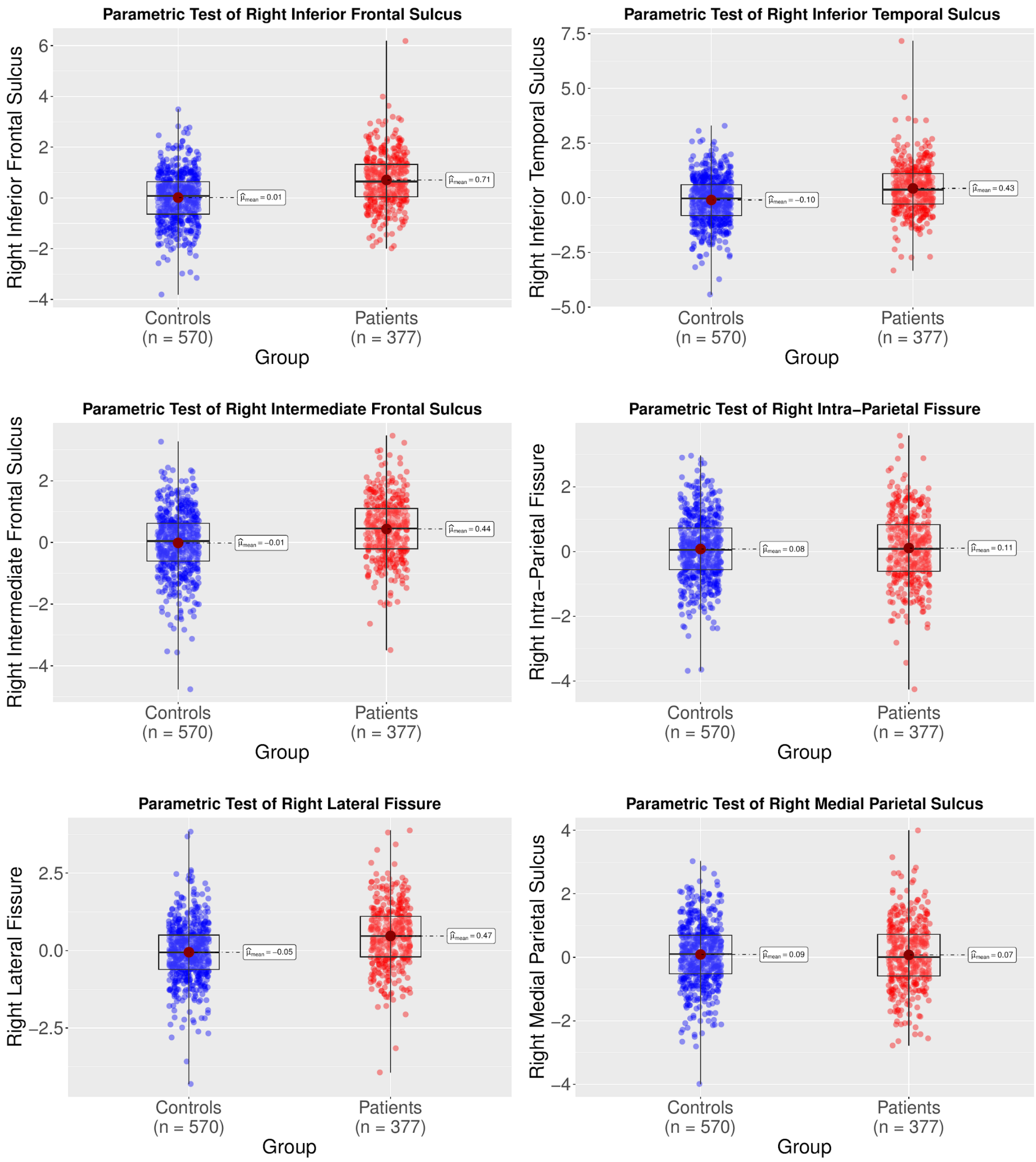

Supplementary Figure 7. Continued.

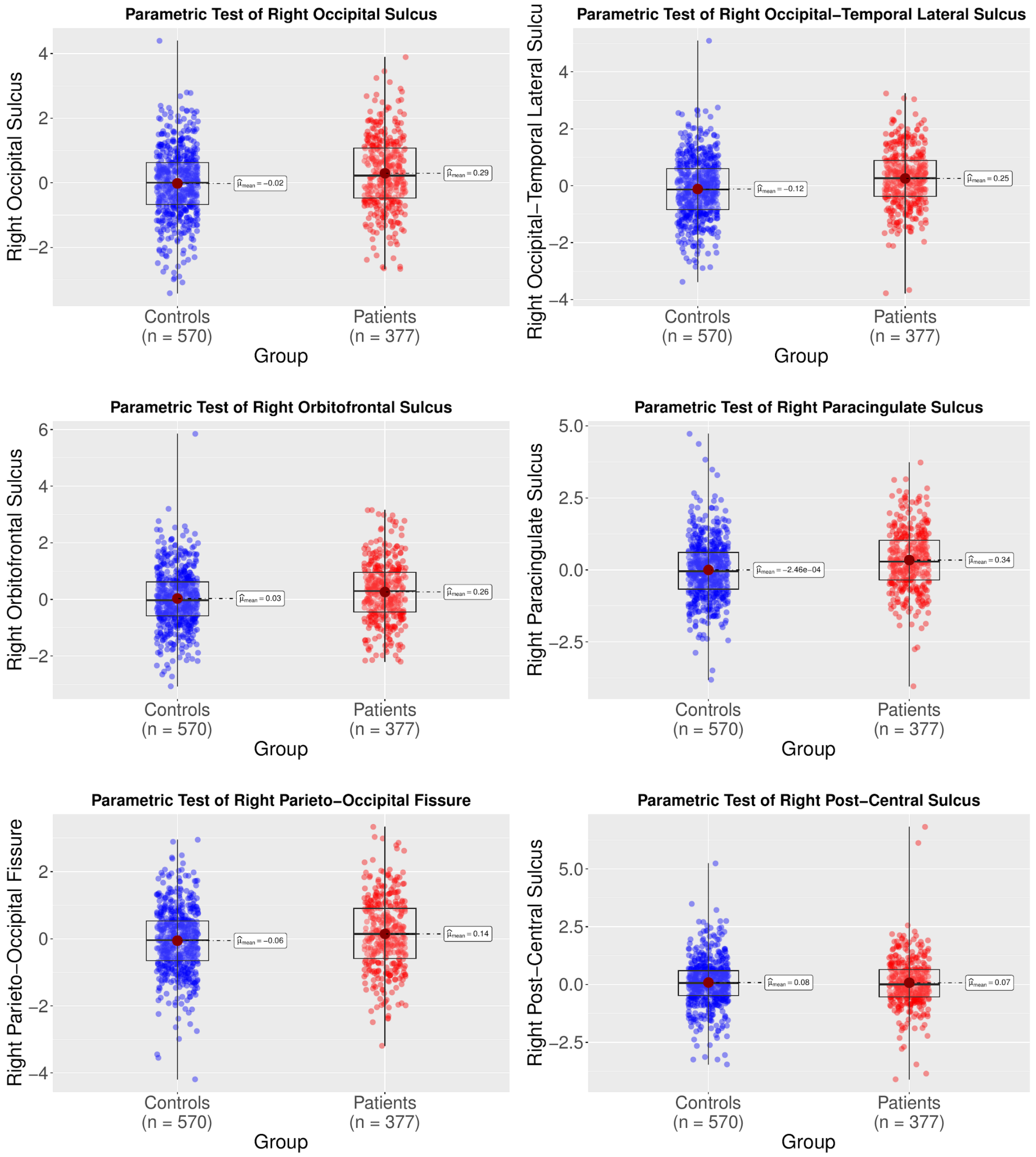

Supplementary Figure 7. Continued.

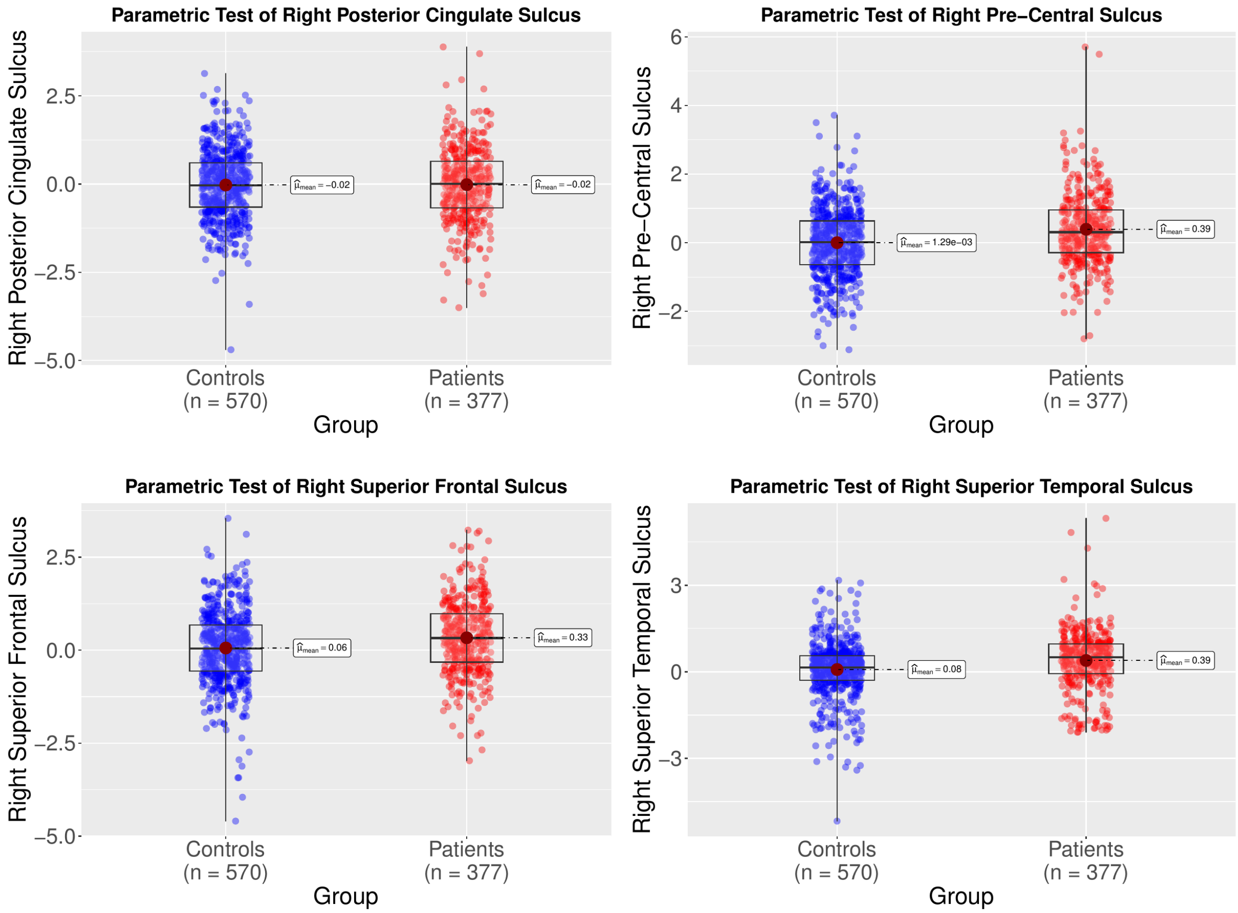

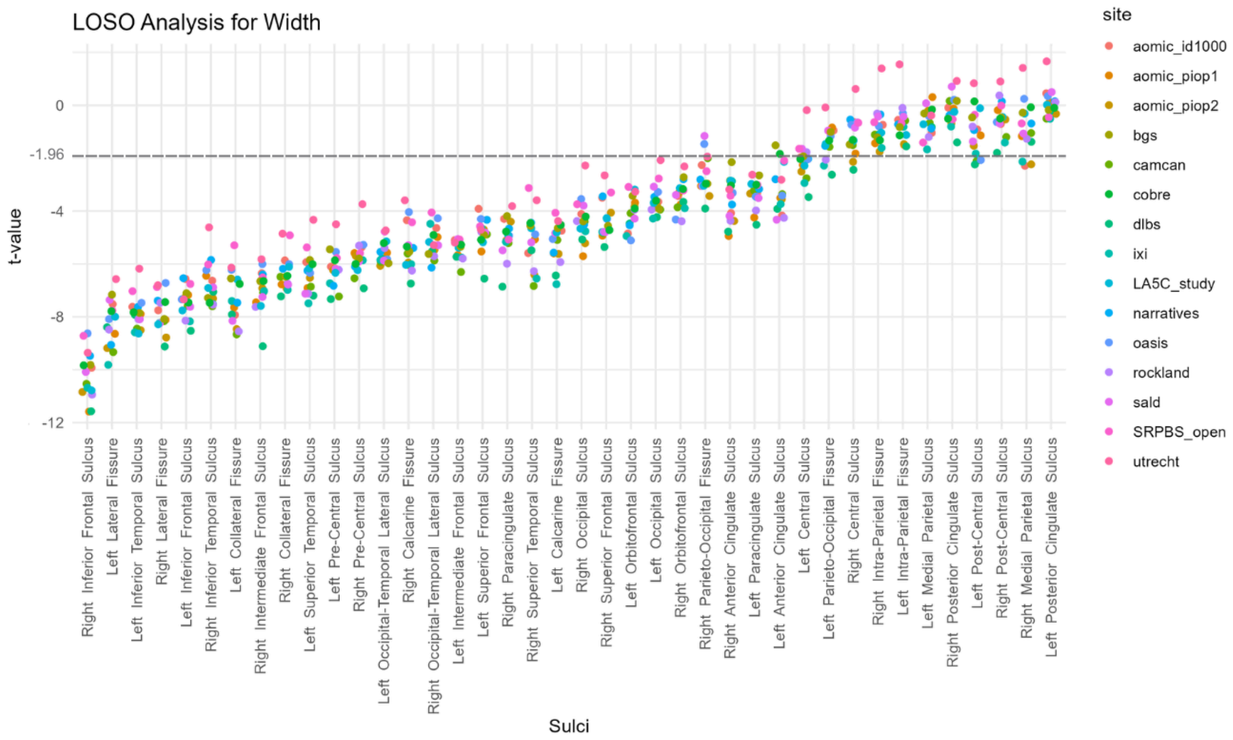


Supplementary Figure 8. We conducted a Leave-One-Site-Out analysis to assess the robustness of diagnostic group differences in normative modeling–based z-scores of sulcal width. Normative modeling was repeated for each iteration, excluding one site at a time (totaling *number of sites – 1* iterations). The horizontally striped line at t = -1.96 indicates $p < 0.05$. The results were consistent across iterations, indicating that the observed diagnostic group differences were not driven by any single site.

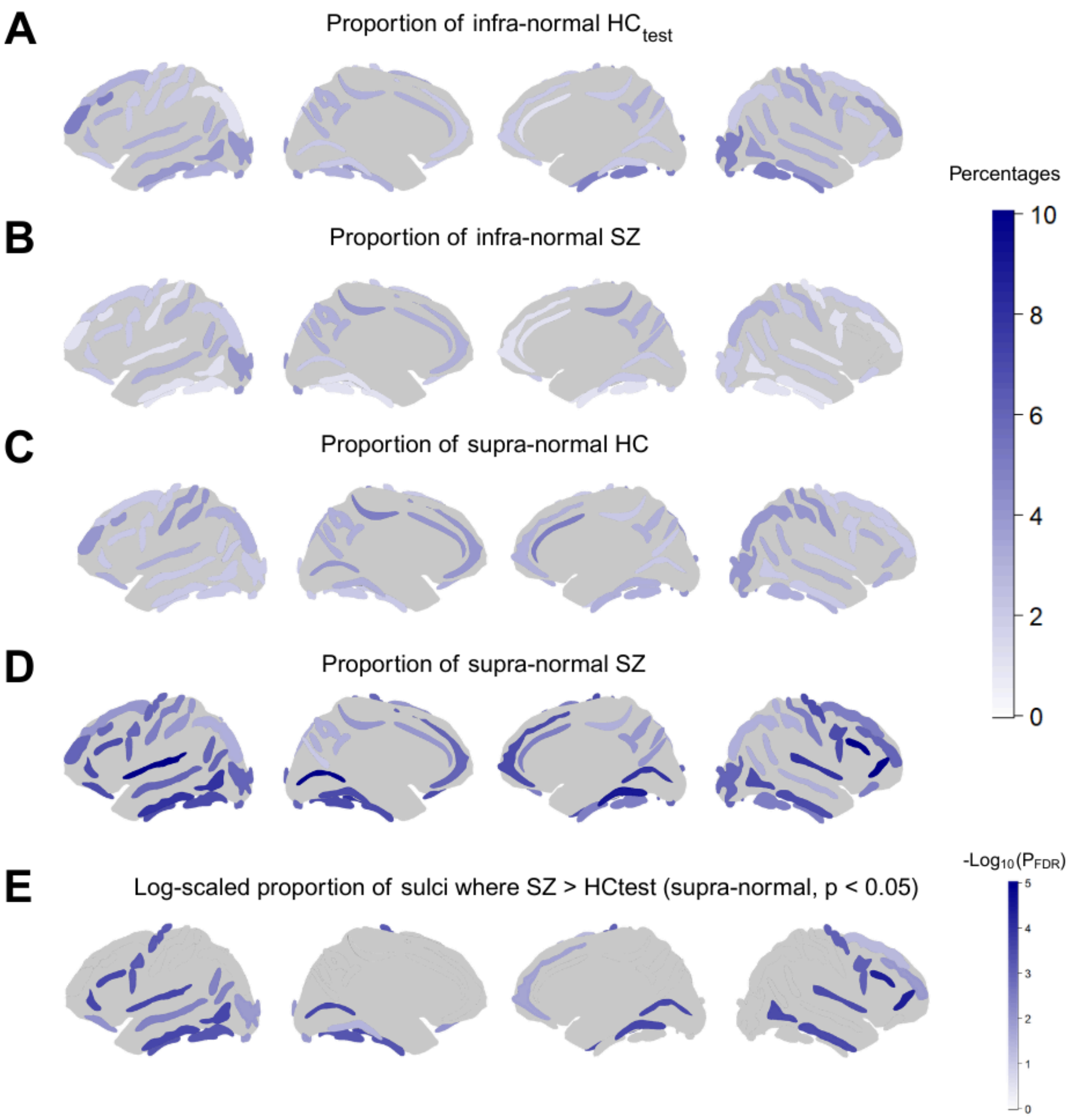


Supplementary Figure 9. Diagnostic group differences in the proportion of individuals with infra- and supra-normal z-scores (defined as z-scores below –1.96 or above 1.96, respectively) between $HC_{test}$ and SZ. Group differences were tested using permuted chi-square tests (n = 10,000) implemented in the *coin* R package, with FDR (q < 0.05) applied for multiple comparisons. (**A**) Proportion of infra-normal z-scores per sulcus in $HC_{test}$. (**B**) Proportion of infra-normal z-scores per sulcus in SZ. (**C**) Proportion of supra-normal z-scores per sulcus in $HC_{test}$. (**D**) Proportion of supra-normal z-scores per sulcus in SZ. (**E**) Sulci showing significant diagnostic group differences in the proportion of supra-normal z-scores, with a greater percentage of SZ individuals exhibiting extreme positive deviations. No significant group differences were observed for infra-normal z-scores.

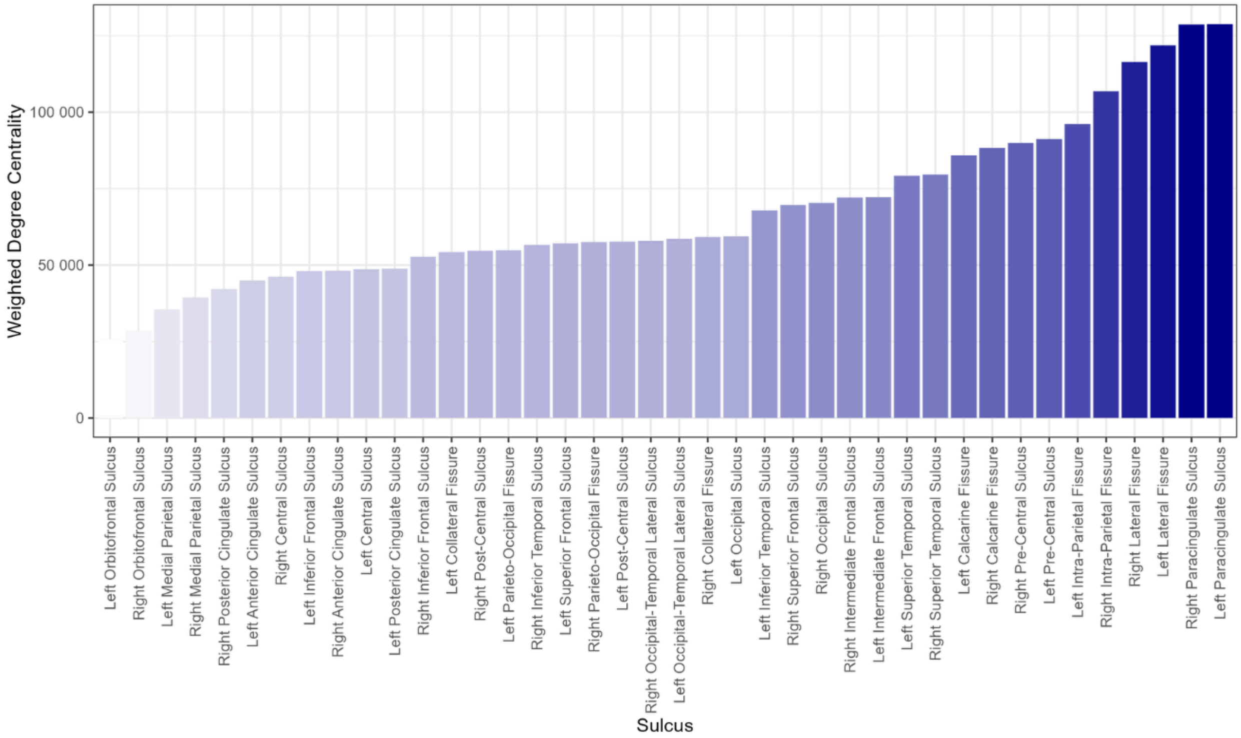


Supplementary Figure 10. Sulci ranked by weighted degree centrality with darker color representing increased weight degree centrality.

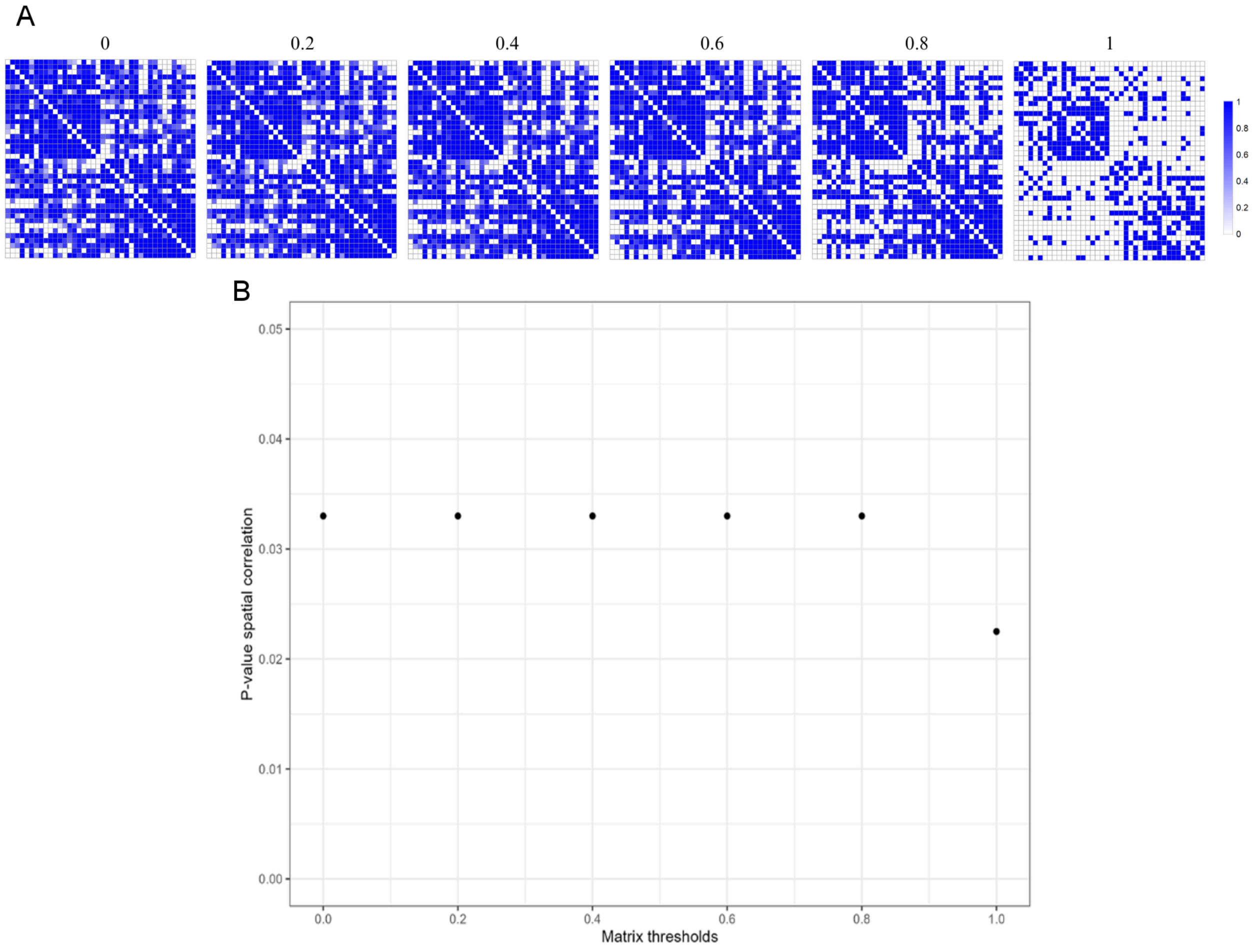


Supplementary Figure 11. **(A)** Consistency matrices visualized across multiple threshold levels. **(B)** *P*-values for the spatial correlations (spin-tests) between reference weighted degree centrality and SZ-related sulcal widening, assessed at varying thresholds of the structural sulcal reference connectivity matrix.

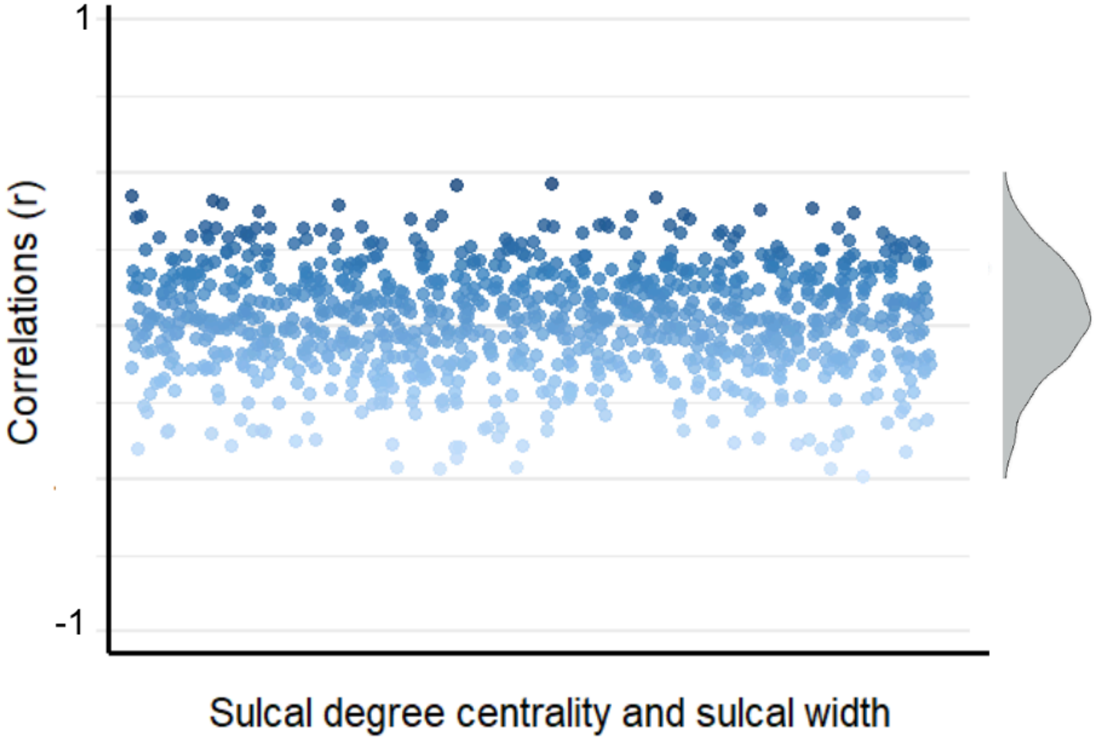


Supplementary Figure 12. Distribution of correlation coefficients of all individuals.

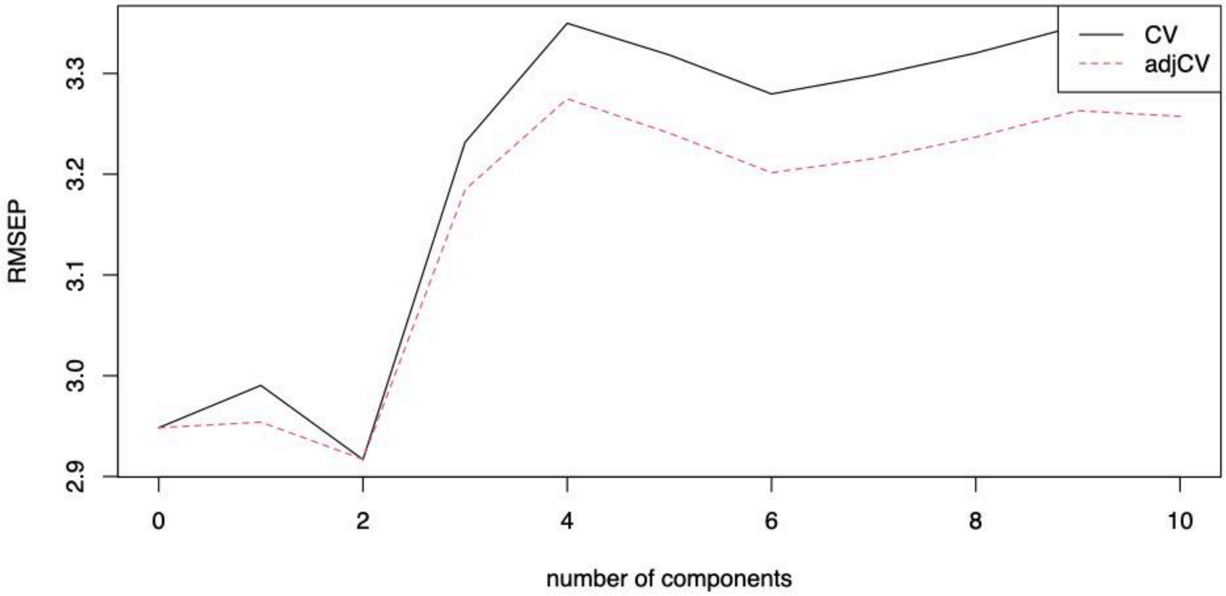


Supplementary Figure 13. Root-mean-squared error of prediction (RMSEP) as a function of the number of latent PLS components. The solid black line represents the raw leave-one-out cross-validation (CV) error, while the dashed red line indicates the bias-corrected CV error (adjCV). PLS, partial least squares.

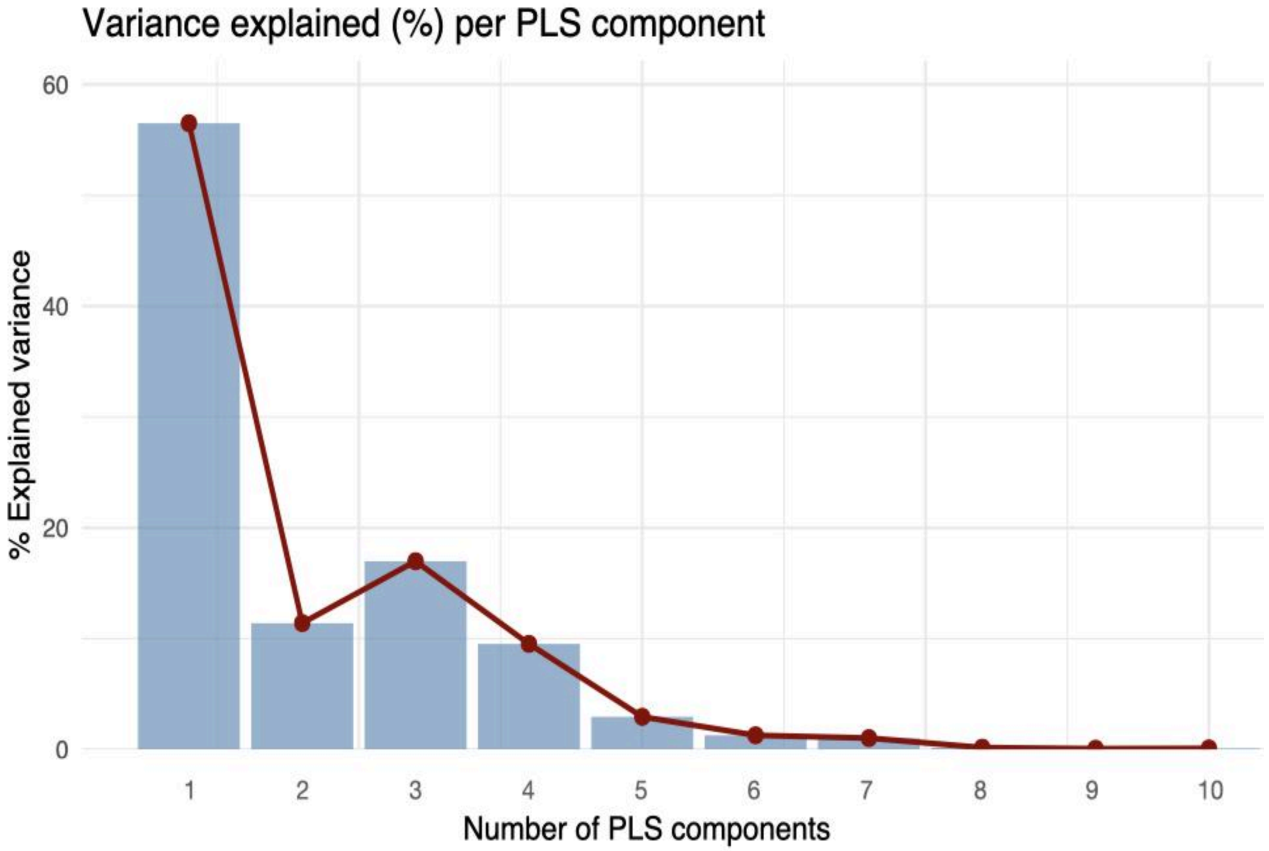


Supplementary Figure 14. Variance explained by the first 10 PLS components. The percentage of variance in diagnostic group differences accounted for by each latent component (PLS1–PLS10) is shown, calculated as the incremental increase in cumulative variance explained. PLS, partial least squares.

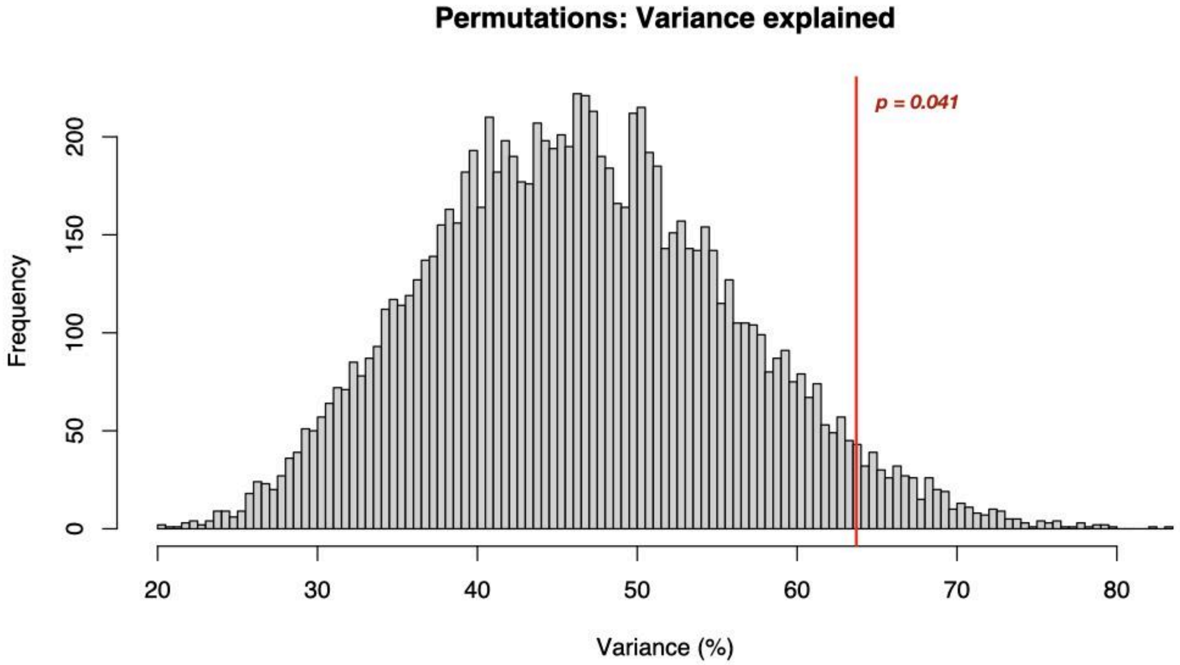


Supplementary Figure 15. Significance of the variance explained by a single-component PLS model (PLS1). The variance explained by the actual PLS1 model (red line) is compared to the null distribution generated from 10,000 PLS models using randomly permuted vectors of the diagnostic group differences. PLS, partial least squares.

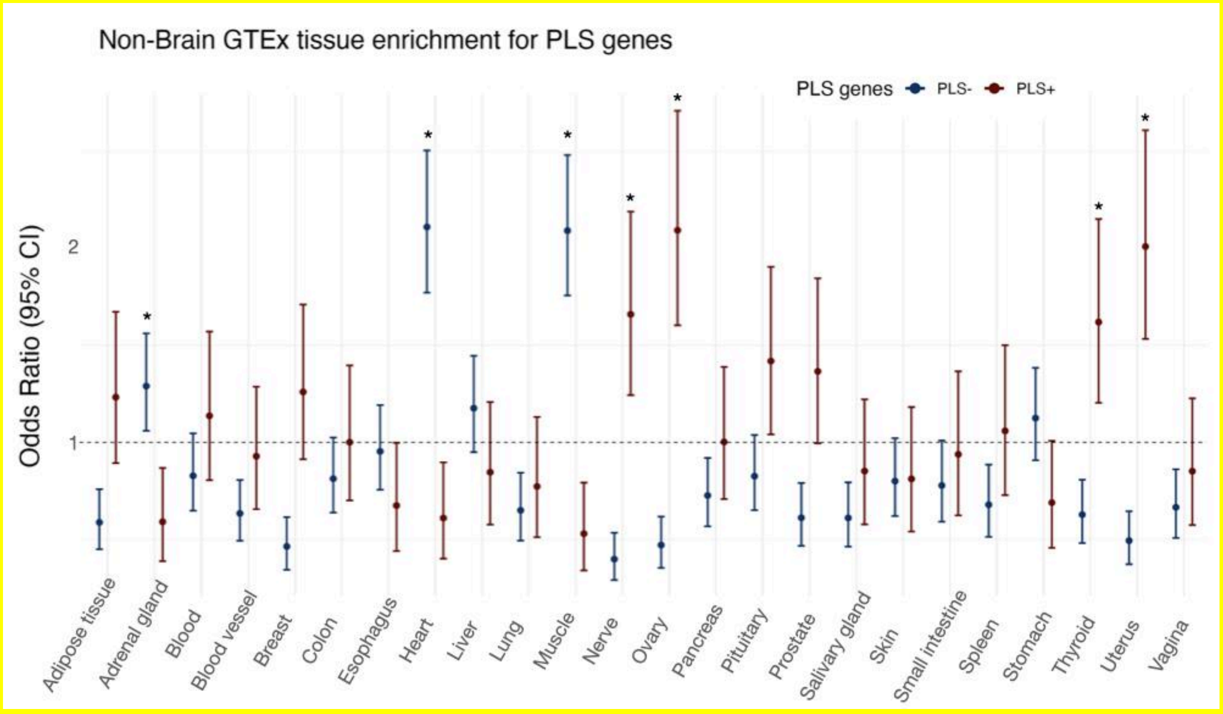


Supplementary Figure 16. Overrepresentation of PLS genes among those specifically expressed in each of the 24 non-brain tissues from GTEx v8 was evaluated using a resampling approach. Specifically, observed overlaps were compared to distributions from 10,000 random gene sets drawn from a background of 15,633 brain-expressed genes. Tissue-specific genes were defined as those within the top 10% of expression specificity scores for each tissue, following established methods (Bryois et al., 2020). Brain regions with overrepresentation of PLS genes across specifically expressed genes were marked with “*”. Error bars indicate 95% confidence intervals of the odds ratios. PLS, partial least squares.

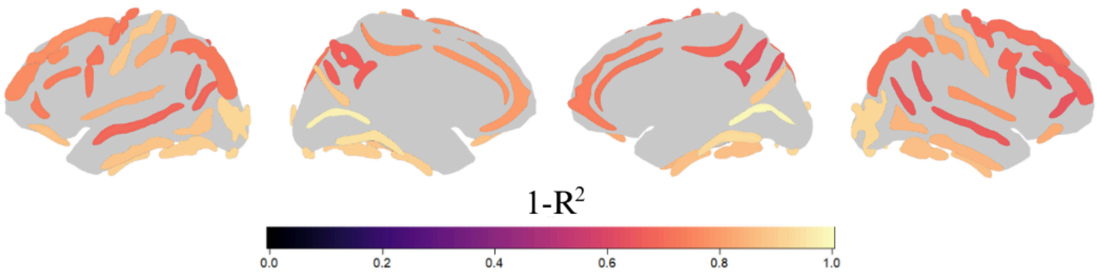


Supplementary Figure 17. The relationship between sulcal width and sulcal thickness. For each sulcus, we applied linear models to predict sulcal width from sulcal thickness and calculated the proportion of unexplained variance ($1 - R^2$). A brighter color indicates higher unexplained variance. For any sulcus the unexplained variance exceeded 64%, indicating that variation in sulcal width is not only driven by changes in sulcal thickness.
